# LINC00599 Promotes Pulmonary Hypertension via LLPS with G3BP1 and MYH9

**DOI:** 10.1101/2024.12.20.629439

**Authors:** Yingqi Wang, Lulu Yin, Shuang Zheng, Aijing Liu, Chunmiao Liu, Zhitu Bao, He Zhu, Xiaoxu Zhao, Ziru Zhao, Yu Pan, Daling Zhu, Hang Yu

## Abstract

**BACKGROUND:** Pulmonary hypertension (PH) represents a significant cardiovascular disorder marked by both functional and structural alterations within the pulmonary vasculature. Long non-coding RNAs (lncRNAs) have been closely associated with the pathogenesis and progression of PH. Nonetheless, the precise mechanisms by which lncRNAs interact with its downstream target molecules to modulate the disease remain inadequately elucidated.

**METHODS:** The expression levels of LINC00599 were quantified in the lung tissues of mice and pulmonary arterial smooth muscle cells (PASMCs) under hypoxic conditions. The involvement of LINC00599 in the progression of PH and vascular remodeling was evaluated through in vivo studies. To investigate the mechanisms by which LINC00599 influences the proliferation of human PASMCs, small interfering RNA and overexpression plasmids were employed.

**RESUITS:** The expression of LINC00599 is upregulated in the medial layer of pulmonary arteries in experimental models of PH and in hypoxic PASMCs. Administration of a single dose of lentivirus-mediated shRNA targeting LINC00599 effectively reverses hypoxic PH in murine models. LINC00599 plays a critical role in PASMC proliferation by modulating stress granule formation through N6-methyladenosine (m6A) modification and promoting proliferation via liquid-liquid phase separation with myosin heavy chain 9. Furthermore, the expression of LINC00599 is regulated by a super-enhancer, which is mediated by the transcription factor ZNF263.

**CONCLUSIONS:** These findings demonstrate for the first time that LINC00599, involved in liquid-liquid phase separation, facilitates the progression of pulmonary hypertension by enhancing the proliferation of pulmonary artery smooth muscle cells.

**GRAPHIC ABSTRACT:** 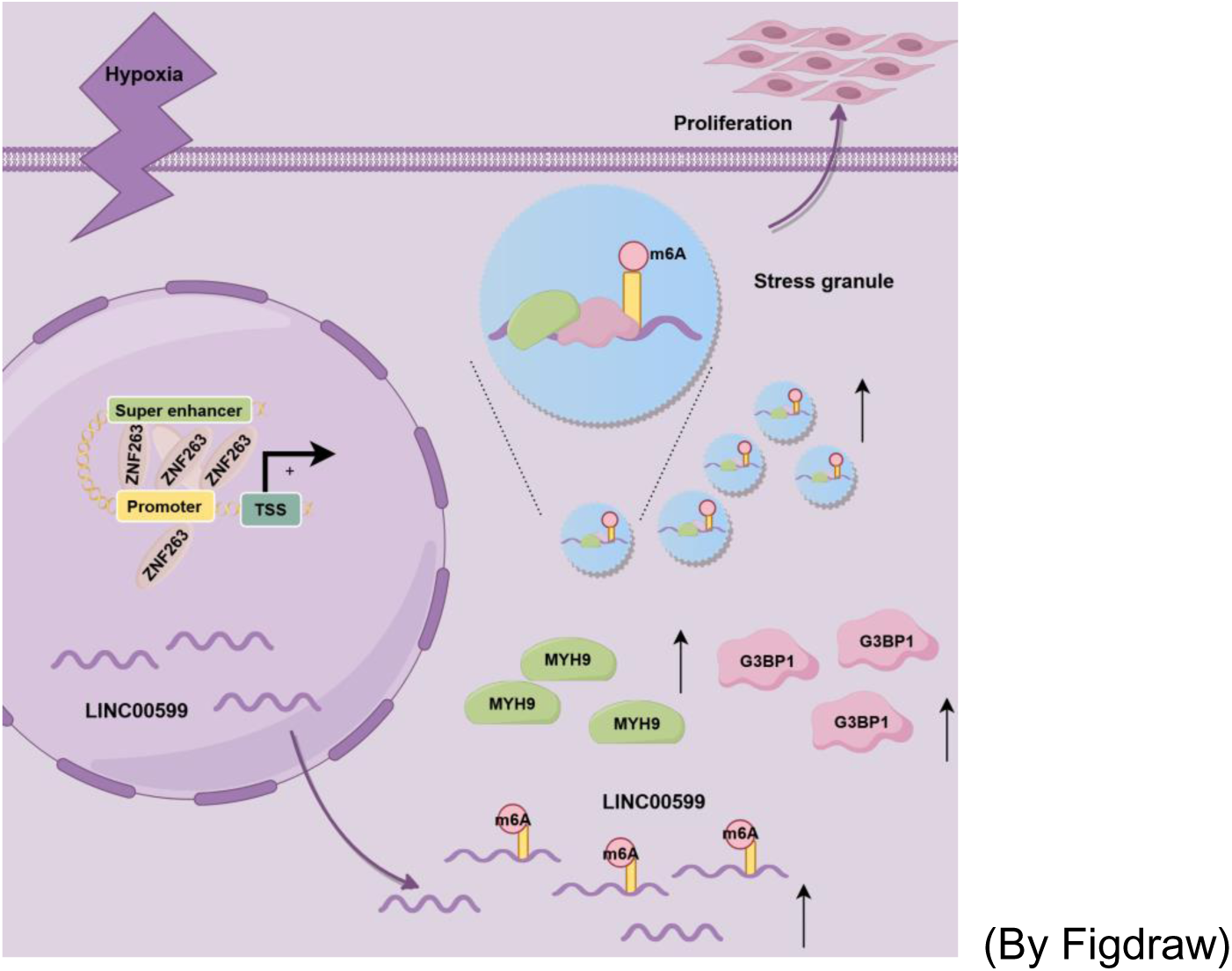

**What Is New?:** This study identified LINC00599 as a critical regulator of liquid-liquid phase separation through m6A modification and as a pivotal target in promoting the progression of PH.

**What Is Relevant?:** LncRNAs are intricately associated with the pathogenesis and progression of PH through the modulation of downstream target molecules. However, it remains unclear whether lncRNAs contribute to PH by forming specific subcellular structures with these target molecules. Furthermore, some studies have suggested that the increased presence of stress granules in PASMCs is implicated in the pathogenesis of pulmonary arterial hypertension. In this study, we provide evidence that LINC00599 promotes the development of PH by forming stress granules with its target proteins. Our findings indicate that the binding of LINC00599, modified with m6A, to G3BP1 and MYH9 in PASMCs suggests that the regulation of liquid-liquid phase separation by lncRNA m6A modification may represent a significant pathological mechanism underlying PH.

**What Are the Pathophysiological Implications?:** We found that LINC00599 levels were elevated in hypoxic PASMCs and experimental PH models. Knockout LINC00599 effectively prevented PH, indicating its potential as both a bomarker and therapeutic target for PH.

## Introduction

Pulmonary hypertension (PH) is a severe cardiovascular disorder marked by both functional and structural alterations in the pulmonary vasculature. These changes result in elevated pulmonary vascular resistance and remodeling, right ventricular hypertrophy, and ultimately, mortality^1–3^. The aberrant proliferation of smooth muscle cells induces excessive collagen synthesis, contributing to arterial wall thickening and luminal narrowing, which collectively drive the pathological remodeling of the pulmonary arteries^4^. Despite substantial progress, the complex mechanisms underlying this process are not yet fully elucidated. A comprehensive understanding of these mechanisms is essential for the development of effective therapeutic strategies for pulmonary hypertension.

Long non-coding RNAs (lncRNAs), which are non-coding RNAs exceeding 200 nucleotides in length, are transcribed by RNA polymerase II. These lncRNAs are involved in a myriad of biological processes, including the regulation of the cell cycle and cellular metabolism through mechanisms such as epigenetic modulation, transcriptional regulation, and post-transcriptional regulation of gene expression^5,6^. Evidence suggests that lncRNAs are intricately linked to the pathogenesis and progression of human diseases by modulating downstream target molecules. This includes mechanisms such as miRNA sponging^7–9^, protein binding^10^, influencing the expression or stability of target proteins^11–13^. Despite this, the mechanisms by which lncRNAs interact with downstream target molecules to regulate disease onset and progression remain inadequately elucidated. Furthermore, the structural composition and morphology of lncRNAs when bound to molecular complexes are still not well understood.

Liquid-liquid phase separation (LLPS) represents a distinctive form of membraneless organelle formation, wherein biomacromolecules aggregate within the cell. This phenomenon is posited to elucidate numerous unresolved questions about cellular mechanisms. In eukaryotic cells, various membraneless structures are demonstrated to arise through LLPS, executing essential functions^14,15^. Stress granules (SGs), which are membraneless organelles assembled via LLPS in the cytoplasm, exemplify this process^16^. Ras GTPase-activating protein-binding proteins 1 and 2 (G3BP1 and G3BP2, respectively) are widely recognized as core components of SGs^17^.

Recent studies have identified lncRNAs within phase separation bodies^18–20^, where they influence disease development and progression by modulating LLPS^21–23^. This discovery establishes a connection between LLPS and lncRNAs, thereby providing a novel perspective on the subcellular localization and functional roles of lncRNAs^24^. Consequently, it is plausible that lncRNAs contribute to the pathological processes of hypoxic pulmonary hypertension (HPH) by regulating SG formation.

N6-Methyladenosine (m6A) is a modification that occurs at the 6th nitrogen atom of adenosine and represents one of the most prevalent modifications in RNA epigenetics^25,26^. The levels of RNA m6A modification are primarily regulated by m6A “writer” proteins, such as METTL3 and METTL14, and “eraser” proteins, including FTO and ALKBH5^27,28^. The m6A modification of lncRNAs serves various functions, such as altering lncRNA structure, mediating transcriptional regulation, participating in mRNA splicing, and regulating lncRNA stability and translation^25^. It has been established that m6A methylation promotes the phase separation of mRNAs^29,30^. Additionally, m6A-modified enhancer RNA facilitates the formation of transcriptional condensates and subsequent gene activation^31^. However, the role of m6A-modified lncRNAs in regulating LLPS and mediating cellular proliferation remains unclear and requires further investigation.

In the current study, we demonstrate that LINC00599 regulates SG formation via m6A modification and affects the proliferation of pulmonary artery smooth muscle cells (PASMCs) under hypoxic conditions. These findings offer morphological evidence elucidating the mechanism underlying hypoxia-induced PASMC proliferation in the context of pulmonary hypertension.

## Results

### Hypoxia Promotes the Expression of LINC00599 in PASMCs and in the Pulmonary Arteries of SuHx mice

The LncBook 2.0 database (https://ngdc.cncb.ac.cn/lncbook) and the LncRNADisease v2.0 database (http://www.rnanut.net/lncrnadisease/) were initially employed to identify the PH-related lncRNA LINC00599 (Fig. 1A). Reverse transcription and real-time PCR (qRT-PCR) assays revealed a significantly elevated expression of LINC00599 in lung tissues of SuHx mouse models (Fig. 1B) and in human pulmonary artery smooth muscle cells (hPASMCs) under hypoxic conditions (Fig. 1C). Fluorescence in situ hybridization (FISH) assays confirmed that LINC00599 was localized in both the cytoplasm and the nucleus of hPASMCs, with hypoxia predominantly increasing the cytoplasmic levels of LINC00599 in hPASMCs exposed to hypoxic conditions (Fig. 1D-E). We also investigated the expression levels of LINC00599 in various mouse tissues and pulmonary artery endothelial cells (PAECs). Our findings indicated that there were no significant differences in LINC00599 expression in the heart, liver, spleen, and kidney tissues of SuHx mouse models (Fig. 1B), as well as in PAECs under hypoxic conditions (Fig. 1F). The expression levels of LINC00599 were significantly elevated in the distal pulmonary arteries of SuHx mice compared to those observed in normoxic mice. (Figure 1G).

**Figure 1.**
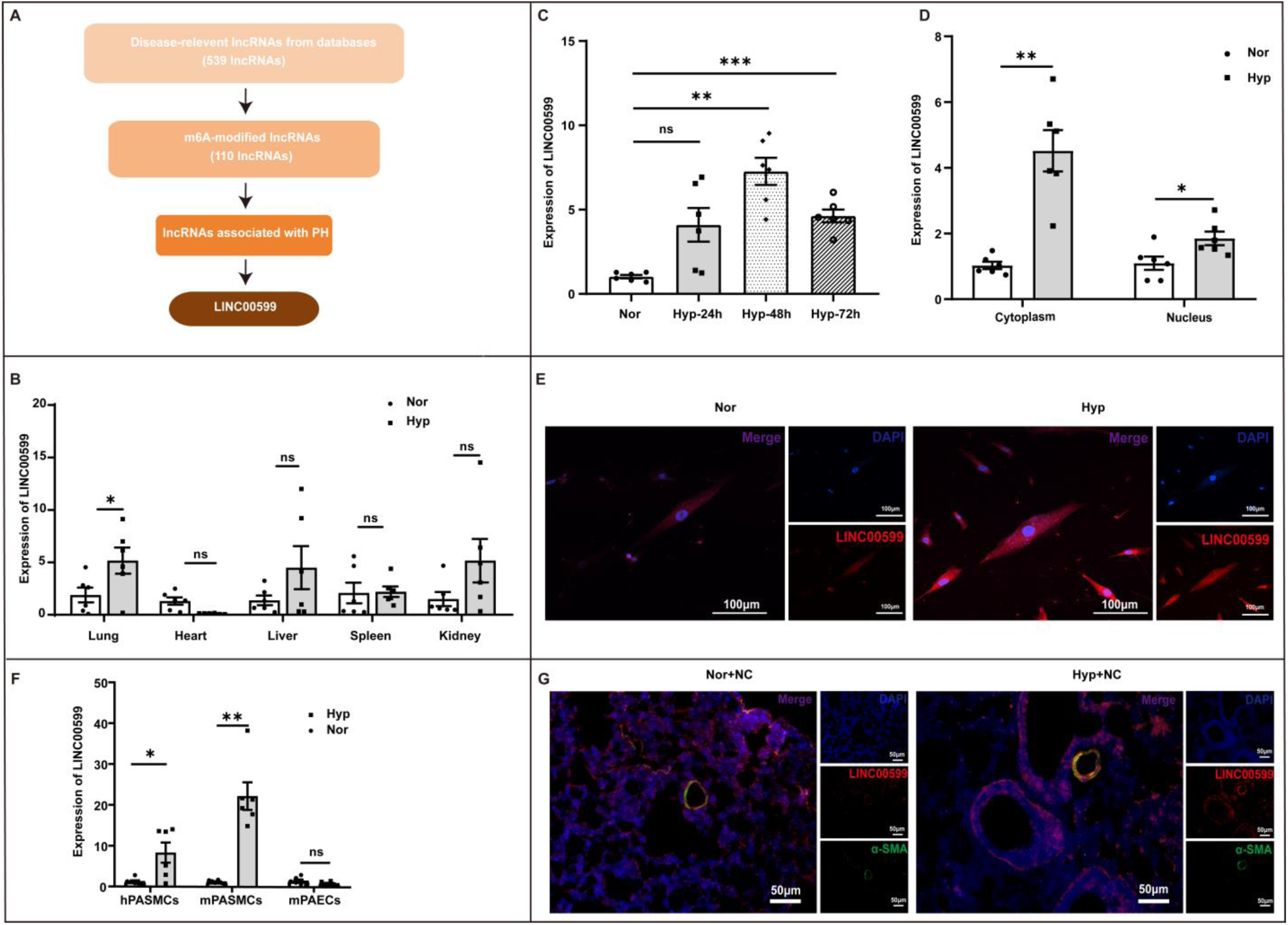
Hypoxia Promotes the Expression of LINC00599 in PASMCs and in the Pulmonary Arteries of SuHx mice. (A)The screening process of LncRNA. (B) qRT-PCR showing the expression of LINC00599 in different tissues of SuHx mouse models (n=6). (C) qRT-PCR analysis of LINC00599 in human PASMCs (n=6). (D) qRT-PCR showing upregulation of LINC00599 in the nucleus and cytoplasm of human PASMCs under hypoxia conditions (n=6). (E) FISH analysis showing upregulation of LINC00599 in the nucleus and cytoplasm of human PASMCs exposed to hypoxia (n=3). Scale bar:100μm. (F) qRT-PCR showing the expression of LINC00599 in PASMCs and PAECs (n=6). (G) FISH and immunofluorescence assay showing the co-location of LINC00599 (red) and α-SMA (green) (n=3). Scale bar:50μm. Data are expressed as the mean±SEM. *\*, P<0.05; **, P<0.01; ***, P<0.001*. ns: no significant; Nor: normoxia; Hyp: hypoxia; NC: negative control; mPASMC: mouse PASMC; mPAEC: mouse PAEC.

### LINC00599-Me-mut Fails to Reverse the Pulmonary Hypertension Amelioration Achieved by LINC00599 shRNA in SuHx Mice

To investigate the potential role of LINC00599 in PH, we demonstrated that the deletion of LINC00599 alleviated SuHx-induced PH. Mice were subjected to a hypoxia chamber for three weeks, followed by administration of a lentivirus carrying LINC00599 shRNA plasmid (sh-LINC00599) or a negative control plasmid (NC) via nasal inhalation. Subsequently, the mice were maintained under normoxic conditions for an additional two weeks (Fig. 2A). The remodeling of distal pulmonary arteries was ameliorated by sh-LINC00599 in SuHx mice (Fig. 2B). Furthermore, under hypoxic conditions, sh-LINC00599 mice demonstrated significant decreases in right ventricular systolic pressure (RVSP) and the ratio of right ventricular to left ventricular plus septum weight (RV/LV+S) compared to NC mice (Fig. 2C, D). Echocardiographic analysis further demonstrated that sh-LINC00599 mice showed enhanced pulmonary arterial velocity time integral (PAVTI) and pulmonary artery acceleration time (PAT) under hypoxic conditions, but not under normoxic conditions (Fig. 2D). The data suggest that the targeted deletion of LINC00599 ameliorates SuHx-induced PH.

**Figure 2.**
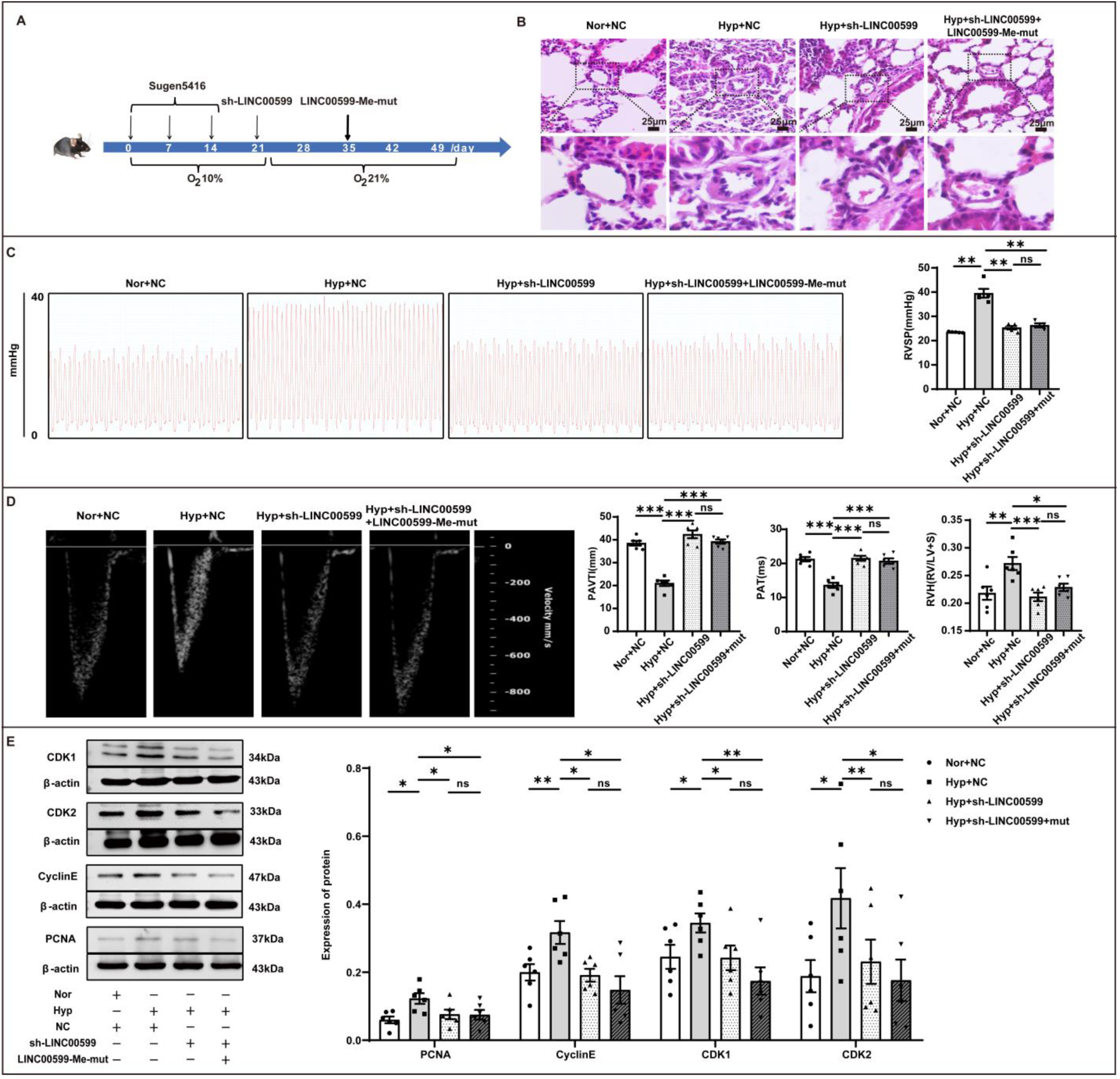
LINC00599-Me-mut Fails to Reverse the Pulmonary Hypertension Amelioration Achieved by LINC00599 shRNA in SuHx Mice. (A) Establishment of SuHx PH mouse models. (B) H&E staining showing the morphology of the pulmonary arteries (n=3). Scale bar: 25μm. (C-D) Echocardiographic measurements and right ventricular catheterization showing the value of pulmonary artery acceleration time (PAT), pulmonary artery velocity time integral (PAVTI), right ventricular (RV)/ left ventricular (LV)+ S weight ratios and right ventricular systolic pressure (RVSP) (n=6). (E) Western blot imagines showing the expression of proliferation-related proteins (n=6). The data are expressed as the mean ± SEM. *\*, P<0.05; **, P<0.01; ***, P<0.001*. ns, no significance; Nor, normoxia; Hyp, hypoxia; NC, negative control; sh-LINC00599: lentivirus carrying LINC00599 shRNA; mut or LINC00599-Me-mut: lentivirus carrying LINC00599 with m6A sites mutation.

The database of RNA modification (RMVar 2.0) showed that m6A modification of LINC00599 was associated with pulmonary hypertension. To investigate the potential role of m6A modification of LINC00599 in PH, we administered a lentivirus carrying LINC00599 plasmids with mutations at the m6A sites (LINC00599-Me-mut) to mice with sh-LINC00599 for an additional two weeks under normoxic conditions (Fig. 2A, S1). However, the values of RVSP, RV/LV+S, PAVTI, and PAT did not exhibit significant changes (Fig. 2C, D). Furthermore, Western blot analysis demonstrated that LINC00599-Me-mut did not exacerbate the levels of proliferation-associated proteins in the lung tissues of SuHx mice, which were ameliorated by sh-LINC00599. (Fig. 2E). These findings suggest that m6A modification of LINC00599 plays a crucial role in PH, potentially due to its effects on the proliferation of PASMCs.

### LINC00599 Regulates the Proliferation of PASMCs

To elucidate the critical role of LINC00599 in the pathogenesis of pulmonary hypertension (PH), we initially employed KEGG enrichment analysis to predict the functional implications of LINC00599. The analysis indicated that LINC00599 is primarily involved in pathways related to cell proliferation (Fig. S2A). Subsequently, we constructed small interfering RNA (si-LINC00599) and overexpression plasmids targeting LINC00599 (Fig. S2B, C). Flow cytometry analysis demonstrated that silencing LINC00599 inhibited the transition from the G1 to the S phase (Fig. 3A) and reduced the viability of hypoxia-exposed human pulmonary artery smooth muscle cells (hPASMCs) compared to cells treated with negative control (NC) siRNA (Fig. 3B). Si-LINC00599 treatment led to decreased levels of CDK1, CDK2, Cyclin A, Cyclin E, and PCNA in hPASMCs (Fig. 3C). The 5-ethynyl-2’-deoxyuridine (EdU) and Ki-67 staining assay demonstrated a reduced proportion of proliferating cells in si-LINC00599-treated cells compared to NC siRNA-treated cells (Fig. 3D, E). These findings suggest that LINC00599 facilitates the proliferation of hPASMCs under hypoxic conditions.

**Figure 3.**
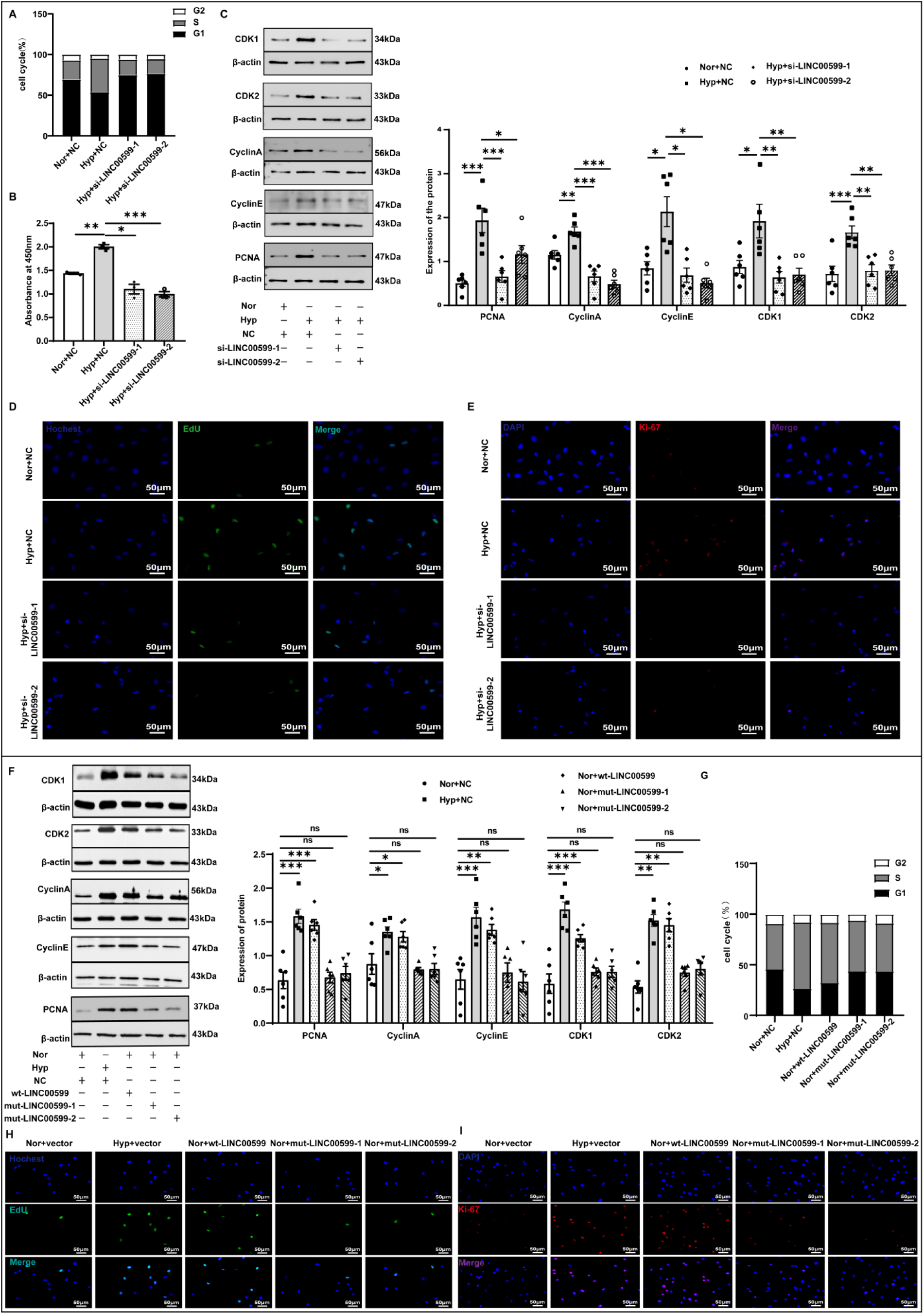
LINC00599 Regulates the Proliferation of PASMCs. (A) Flow cytometry showing the changes of cell cycle in hypoxic hPASMCs treatment with LINC00599 siRNA (n=6). (B) CCK-8 assay showing the cell viability of hPASMCs (n=3). (C) Western blot assay showing the levels of proliferation-relevant proteins in hPASMCs exposed to hypoxia treated with LINC00599 siRNA (n=6). (D) EdU assay and (E) Ki-67 immunofluorescence showing the proliferation of hPASMCs under hypoxia treated with LINC00599 siRNA (n=3). Scale bar: 50μm. (F) Flow cytometry showing the changes of cell cycle in normaxic hPASMCs treatment with LINC00599 plasmids (n=6). (G) Western blot assay showing the levels of proliferation-relevant proteins in normaxic hPASMCs treatment with LINC00599 plasmids (n=6). (H) EdU assay and (I) Ki-67 immunofluorescence showing the proliferation of hPASMCs under normoxia treated with LINC00599 plasmids (n=3). Scale bar: 50μm. Data are expressed as mean ±SEM. *\*\*, P<0.01; ***, P<0.001*; Nor: normoxia; Hyp: hypoxia; si-LINC00599: LINC00599 siRNA; NC: negative control; wt-LINC00599:

To further investigate the role of elevated LINC00599 expression in hPASMCs, we overexpressed LINC00599 in normoxic hPASMCs using plasmids encoding the wild-type LINC00599 (wt-LINC00599). Constitutive expression of LINC00599 facilitated the transition from the G1 to S phase (Fig. 3F) and upregulated the levels of CDK1, CDK2, Cyclin A, Cyclin E, andPCNA Ki-67 (Fig. 3G). Additionally, EdU and Ki-67 staining assay indicated a higher proportion of proliferating cells in wt-LINC00599-treated cells compared to those treated with negative control (NC) plasmids (Fig. 3H, I).

### LINC00599 Regulates Hypoxia-Induced LLPS through m6A Modification and Mediates PASMC Proliferation

To investigate the mechanism by which LINC00599 influences cellular proliferation, we initially employed bioinformatics analysis to predict the m6A methylation sites on LINC00599 (Fig. 4A). RNA immunoprecipitation (RIP) assay demonstrated that the m6A levels of LINC00599 were elevated in hPASMCs under hypoxic conditions compared to normoxic conditions (Fig. 4B). To elucidate the mechanism underlying hypoxia-induced m6A modification of LINC00599, we assessed the expression levels of the m6A methyltransferase METTL3 and the demethylase ALKBH5. Our data indicated that hypoxia upregulated METTL3 expression while downregulating ALKBH5, resulting in an elevated m6A level of LINC00599 in hPASMCs (Fig. 4C). To investigate the downstream mechanisms by which LINC00599 influences hPASMC proliferation, we conducted a Gene Ontology (GO) analysis to predict its biological functions. The analysis revealed an association between LINC00599 and cytoplasmic stress granules (Fig. S3A). we have shown that elevated levels of LINC00599 are primarily localized within the cytoplasm of hPASMCs (Fig. 1D, E). Consequently, we hypothesize that LINC00599 regulates the formation of stress granules through m6A modification. Bioinformatics analyses predicted the interacting proteins of LINC00599, indicating a potential interaction between LINC00599 and G3BP1 (Fig. 4D, S3B). Subsequent molecular docking analysis identified the specific interaction sites between LINC00599 and G3BP1 (Fig. S3C). The binding of LINC00599 to G3BP1 was experimentally validated through RNA pull-down and RNA immunoprecipitation (RIP) assays (Fig. 4E, F). Furthermore, fluorescence in situ hybridization (FISH) and immunofluorescence assays demonstrated the co-localization of LINC00599 and G3BP1 in hypoxia-exposed hPASMCs, which was disrupted by the phase separation inhibitor 1,6-hexanediol (Fig. 4G, H). Silencing of LINC00599 decreased the interaction between LINC00599 and G3BP1, leading to a reduction in cytoplasmic condensates in hPASMCs (Fig. 4G). This observation suggests that the binding between LINC00599 and G3BP1 exhibits characteristics of LLPS. Notably, the reduction in condensates induced by si-LINC00599 could be reversed by co-transfection with wild-type LINC00599 plasmids (wt-LINC00599), but not with plasmids containing mutations at m6A sites (mut-LINC00599) (Fig. 4H). This indicates that LINC00599 regulates LLPS through m6A modification.

**Figure 4.**
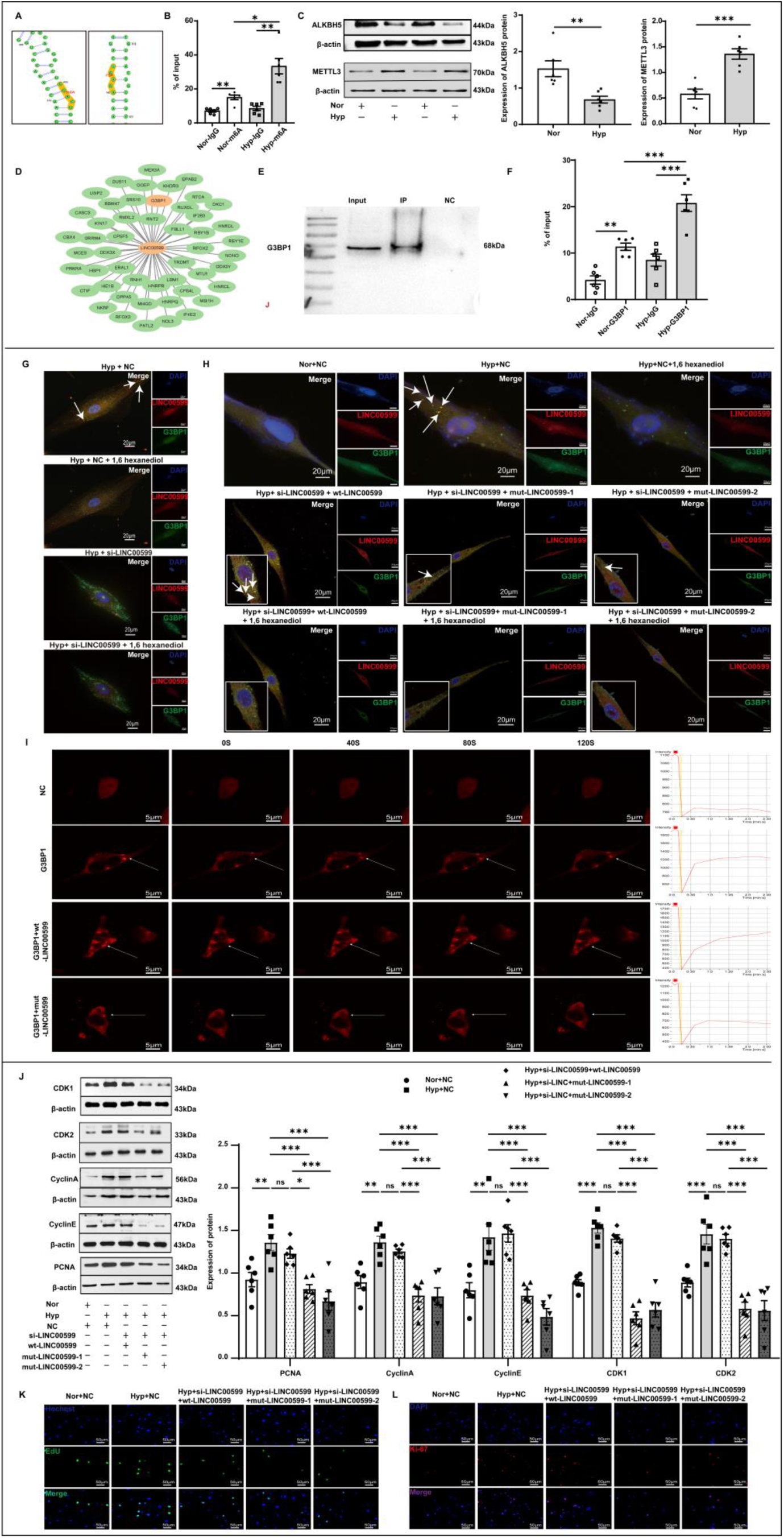
LINC00599 regulates hypoxia-induced LLPS through m6A modification and mediates PASMC proliferation. (A) Bioinformatic analysis predicting m6A sites on LINC00599. (B) RIP assay results indicating m6A modification levels of LINC00599 (n=6). (C) Western blot images depicting protein levels of ALKBH5 and METTL3 in hPASMCs (n=6). (D) Bioinformatic prediction of proteins interacting with LINC00599. (E-F) RNA pull-down and RIP assays demonstrating the interaction between LINC00599 and G3BP1 (n=6). (G) FISH and immunofluorescence assays illustrating the interaction between LINC00599 and G3BP1 following treatment with 1,6-hexanediol in hPASMCs (n=3). Scale bar: 20μm. (H) FISH-immunofluorescence assay showing the co-localization of LINC00599 and G3BP1 (n=3). Scale bar: 20μm. (I) FRAP assay demonstrating the recovery of bleached puncta (n=3). Scale bar: 5μm. (J) Western blot analysis illustrating the expression levels of proteins associated with proliferation (n=6). (K) The EdU assay indicating the proliferation of hPASMCs (n=3). Scale bar: 50μm. (L) Ki-67 immunofluorescence revealing the proliferation of hPASMCs (n=3). Scale bar: 50μm. Data are expressed as mean ±SEM. *\*,P<0.05; **, P<0.01; ***, P<0.001.* Nor, normoxia; Hyp, hypoxia; NC, negative control; wt-LINC00599, overexpressing wild type LINC00599 plasmid; mut-LINC00599, overexpressing LINC00599 plasmid with m6A sites mutation; si-LINC00599,LINC00599 siRNA.

We also investigated the potential phase separation of LINC00599 with G3BP1 in 293T cells by co-transfecting cells with G3BP1 fluorescent plasmids alongside either wt-LINC00599 or mut-LINC00599 plasmids. Utilizing the fluorescence recovery after photobleaching (FRAP) assay, we observed that the bleached puncta exhibited recovery in cells co-transfected with G3BP1 and wt-LINC00599 plasmids, whereas no recovery was detected in cells co-transfected with G3BP1 and mut-LINC00599 plasmids (Fig. 4I). Additionally, the FISH-immunofluorescence assay revealed a higher number of cytoplasmic condensates in 293T cells co-transfected with G3BP1 and wt-LINC00599 plasmids compared to those co-transfected with G3BP1 and mut-LINC00599 plasmids (Fig. S3D). The findings further corroborated that LINC00599, functioning as a crucial component, modulates LLPS in conjunction with G3BP1 through m6A modification.

Subsequently, we conducted additional experiments to determine whether the disruption of LINC00599-G3BP1 phase separation affects the proliferation of hPASMCs under hypoxic conditions. Our findings revealed that silencing LINC00599 (si-LINC00599) attenuated the hypoxia-induced proliferation of hPASMCs. This attenuation could be reversed by co-transfection with wt-LINC00599 plasmids, but not with mut-LINC00599 plasmids (Fig. 4J-L). These results suggest that the disruption of LINC00599-mediated LLPS mitigates the hypoxia-induced proliferation of hPASMCs.

### MYH9 is a Novel Binding Partner of LINC00599 and Mediates Hypoxic PASMC Proliferation via Stress Granules

To investigate the downstream targets by which LINC00599 mediates the proliferation of pulmonary artery smooth muscle cells (PASMCs), proteins interacting with LINC00599 were identified using liquid chromatography-tandem mass spectrometry (LC-MS/MS). Myosin heavy chain 9 (MYH9) emerged as the top candidate and was subsequently selected for further experimentation (Fig. S4A). RNA pull-down and RIP assays confirmed the interaction between LINC00599 and MYH9 (Fig. 5A, B). The Co-immunoprecipitation (Co-IP) assay demonstrated an interaction between MYH9 and G3BP1 (Fig. 5C). Western blot analysis revealed an upregulation of MYH9 in human PASMCs under hypoxic conditions, as well as in the lung tissues of SuHx mouse models (Fig. 5D). These findings imply that LINC00599, MYH9, and G3BP1 form a complex. Bioinformatics analysis predicted the presence of an intrinsically disordered region within the MYH9 protein (Fig. S4B). Fluorescence in situ hybridization (FISH) and immunofluorescence assays further confirmed that the LINC00599-MYH9-G3BP1 complex was disrupted by 1,6-hexanediol treatment (Fig. 5E). Additionally, silencing MYH9 inhibited the formation of stress granules, indicating that MYH9 is a component of stress granules in hPASMCs (Fig. 5E).

**Figure 5.**
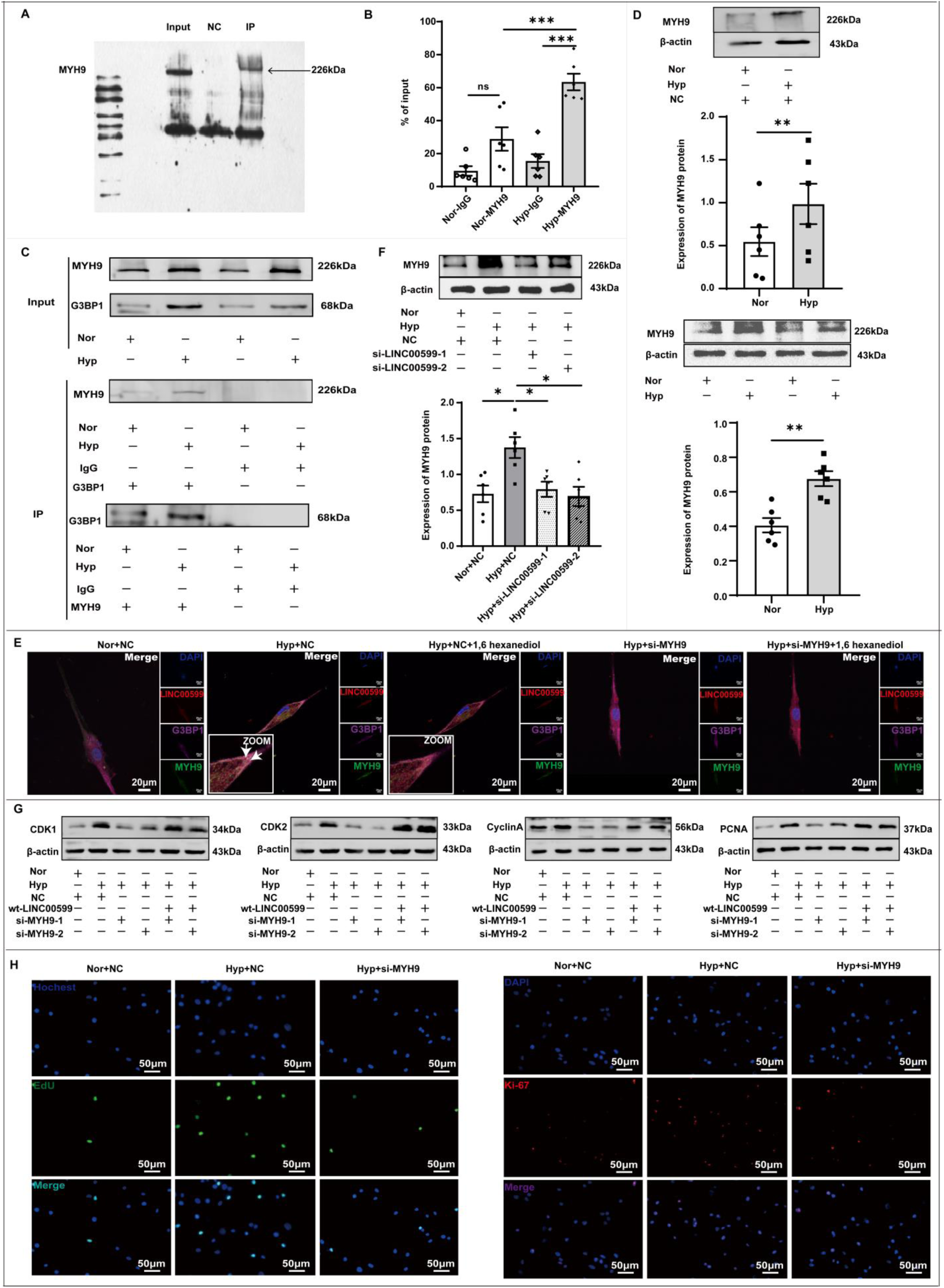
MYH9 is a novel binding partner of LINC00599 and mediates hypoxic PASMC proliferation via stress granules. (A) RNA pull-down and (B) RIP assays showing the interaction between MYH9 and LINC00599 (n=6). (C) Co-immunoprecipitation analysis confirming the binding of G3BP1 to MYH9 (n=3). (D) Western blot analysis assessing the expression levels of MYH9 (n=6). (E) FISH combined with immunofluorescence assays revealing the interaction among LINC00599, G3BP1, and MYH9 (n=3). Scale bar: 20μm.(F) Western blot images illustrating the impact of si-LINC00599 on MYH9 protein levels (n=7). (G) Western blot images depicting the expression of proteins associated with cell proliferation (n=6). (H) EdU and Ki-67 assays demonstrating the effects of MYH9 siRNA on the proliferation of hPASMCs (n=3). Scale bar: 50μm. The results are expressed as mean ± SEM. *\*, P<0.05; **, P<0.01; ***, P<0.001*. ns: no significance; Nor: normoxia; Hyp: hypoxia; NC: negative control; wt-LINC00599: overexpressing wild type LINC00599 plasmid; si-LINC00599: LINC00599 siRNA; si-MYH9: MYH9 siRNA.

To investigate whether LINC00599 regulates MYH9 expression under hypoxic conditions, we treated hPASMCs with siRNA targeting LINC00599 (si-LINC00599). The results indicated that MYH9 levels were significantly decreased when hypoxic cells were transfected with si-LINC00599 concurrently (Fig. 5F). These findings suggest that LINC00599 modulates MYH9 expression in response to hypoxia. Additionally, silencing MYH9 resulted in reduced proliferation of hPASMCs under hypoxic conditions (Fig. 5G, H, S4D, F). This proliferative capacity was restored when cells were co-transfected with LINC00599 wild-type plasmids (Fig. 5G, S4F). The presented data indicate that LINC00599 modulates the proliferation of hPASMCs under hypoxic conditions through the mediation of MYH9.

### Super-Enhancer Regulates LINC00599 Expression through Transcription Factor ZNF263 in PASMCs

To explore the upstream regulatory mechanisms of LINC00599, we identified a super-enhancer, designated as SE-2082886, which comprises six typical enhancers and may play a role in its regulation (Fig. S5A). Additionally, we identified ZNF263 as a potential transcription factor with binding affinity for both the super-enhancer and the promoter regions of LINC00599 (Fig. S5B). To assess the functional role of the SE in the expression of LINC00599, we employed JQ-1, a known SE inhibitor, to evaluate the levels of LINC00599 in hypoxia-exposed hPASMCs. Our findings demonstrated a reduction in LINC00599 expression in response to JQ-1 treatment, indicating a significant correlation between SEs and LINC00599 (Fig. 6A). Additionally, we observed that hypoxia led to an upregulation of ZNF263 in hPASMCs (Fig. 6B). Silencing ZNF263 under hypoxic conditions resulted in decreased LINC00599 expression (Fig. 6C, S5C), thereby confirming the regulatory role of ZNF263 on LINC00599 expression. Furthermore, we explored the upstream mechanisms governing the expression of LINC00599. We partitioned the average promoter region of LINC00599 into three distinct segments. Our findings indicated that the histone activity markers H3K27ac and H3K4me3, along with ZNF263, were enriched in the second segment of the promoter (Fig. 6D, E, S5D). Co-immunoprecipitation (Co-IP) assays demonstrated an interaction between H3K27ac and ZNF263 (Fig. 6F). Furthermore, chromatin immunoprecipitation-quantitative polymerase chain reaction (ChIP-qPCR) assays revealed that ZNF263 binds to the E2 region of the SE-2082886 enhancer (Fig. 6G). These results collectively suggest that ZNF263 transcriptionally regulates the expression of LINC00599 by binding to both the E2 region of SE-2082886 and the second segment of the LINC00599 promoter.

**Figure 6.**
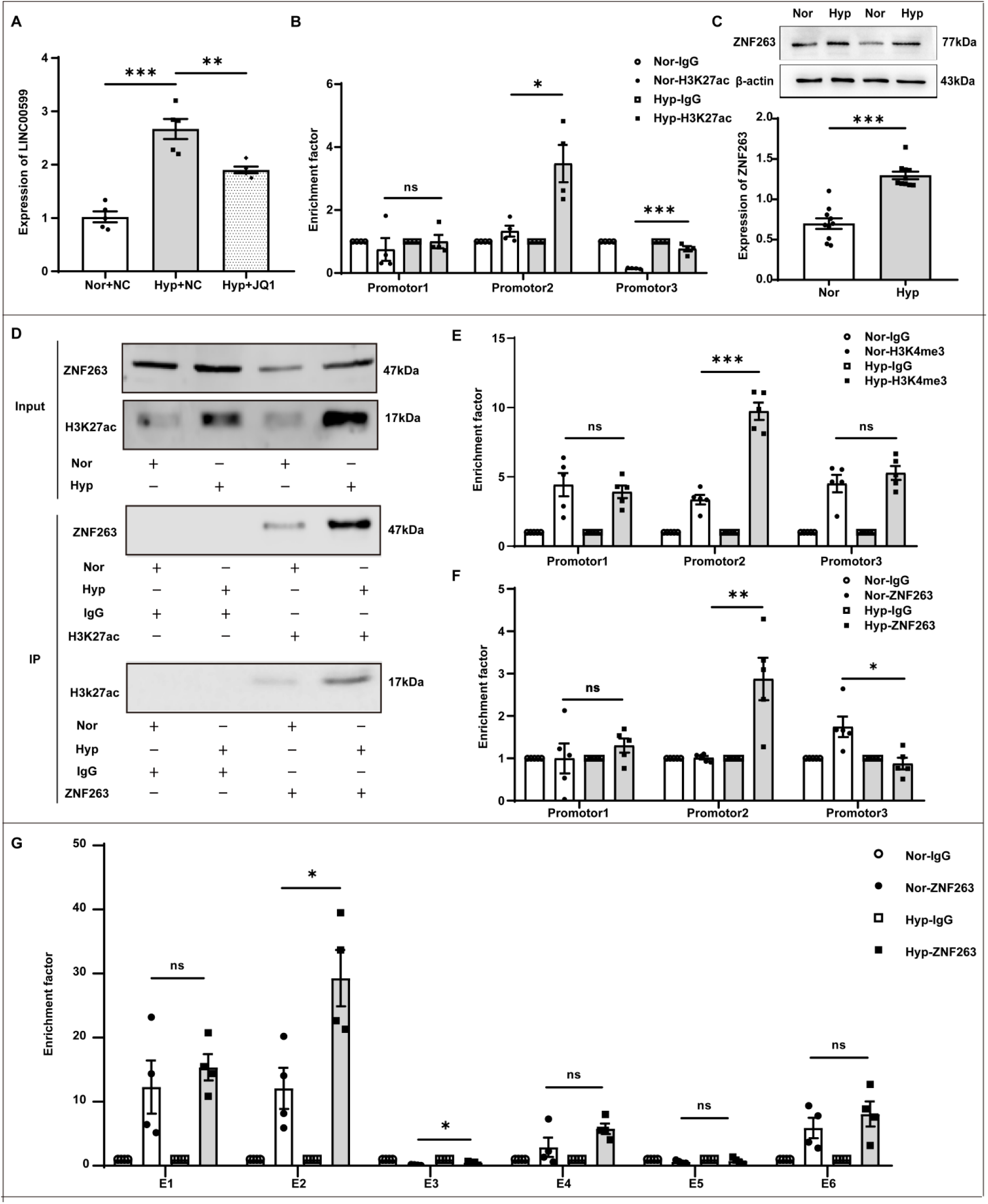
Super-enhancer regulates LINC00599 expression through transcription factor ZNF263 in PASMCs. (A) qPCR analysis demonstrating the expression of LINC00599 (n=5). (B) Western blot analysis evaluating the expression levels of ZNF263 (n=10). (C) qPCR illustrating the effect of si-ZNF263 on LINC00599 expression levels (n=6). (D-E) ChIP-PCR indicating the enrichment of H3K4me3 and ZNF263 at the LINC00599 promoter region (n=5). (F) Co-immunoprecipitation assay demonstrating the interaction between H3K27ac and ZNF263. (G) ChIP-qPCR showing the binding of ZNF263 to the E2 region of the super enhancer (n=4). The data are expressed as the mean ± SEM. *, *P<0.05; **, P<0.01; ***, P<0.001*. ns: no significance; Nor: normoxia; Hyp: hypoxia; NC: negative control; si-ZNF263: ZNF263 siRNA; E: typical enhancer.

## Discussion

LncRNAs play crucial roles in pulmonary hypertension. In this study, we discovered that LINC00599 regulates the proliferation of PASMCs under hypoxic conditions. Mechanistically, LINC00599 modulates stress granule (SG) formation by binding to G3BP1 via m6A modification. Additionally, MYH9 is incorporated into SGs through its interaction with LINC00599 and mediates PASMC proliferation as a downstream target of LINC00599. The upregulation of LINC00599 is controlled by a super enhancer through the transcription factor ZNF263. These findings suggest that LINC00599 could serve as a potential therapeutic target for hypoxia-induced PH.

Growing evidence suggests that lncRNAs are critically involved in the pathological processes of various diseases. For example, lncRNA H19 upregulates the expression of lactate dehydrogenase A (LDHA) by inhibiting microRNA-519d-3p, thereby inducing glycolysis and promoting the proliferation of gastric tumor cells^32^. In contrast, lncDACH1 attenuates lung fibrosis by interacting with serine/arginine-rich splicing factor 1 (SRSF1) to inhibit the accumulation of catenin beta-1 (CTNNB1), highlighting its potential as a therapeutic target for pulmonary fibrosis^33^. These findings underscore the importance of the tissue-specific distribution of lncRNAs in regulating disease pathology. Our data demonstrated that LINC00599 was expressed across multiple tissues; however, hypoxia predominantly up-regulated LINC00599 levels in lung tissue. Subsequent experiments confirmed that the elevated levels of LINC00599 occurred specifically in PASMCs rather than PAECs. These findings indicate that LINC00599 exhibits tissue- and cell-specific distribution. Knockdown of LINC00599 using specific siRNA significantly enhanced the proliferation of PASMCs under hypoxic conditions. Moreover, interference with LINC00599 expression via lentivirus carrying LINC00599 shRNA plasmid (sh-LINC00599) reversed hypoxia-induced PH in vivo. Therefore, the cell-specific distribution characteristics of LINC00599, along with its tissue-specific distribution, may offer potential targets for the development of targeted therapeutics for pulmonary hypertension.

Stress granules (SGs), a special form of liquid-liquid phase separation (LLPS), are large, non-membrane-bound cytoplasmic entities that occur in cells exposed to various stresses, including heat, viral infection, oxidative conditions, ultraviolet irradiation, and hypoxia^34^. It has been reported that lncRNAs are implicated in SG formation. For instance, lncRNA TM4SF1-AS1 facilitates SG formation and inhibits apoptosis in gastric cancer cells by sequestering RACK1, an activator of the stress-responsive MAPK pathway, within SGs^35^. However, the mechanisms through which lncRNAs interact with other molecules in SGs, the specific proteins to which lncRNAs bind, and the details of these interactions remain unclear. In the present study, we employed bioinformatics analysis to predict proteins that interact with LINC00599 and identified G3BP1, a protein typically associated with SGs, as a potential binding candidate. Given that lncRNAs have been validated to facilitate liquid-liquid phase separation of proteins^21,36,37^, we subsequently investigated the potential interaction between LINC00599 and the G3BP1 protein through fluorescence co-localization. Our findings revealed that the interaction between LINC00599 and G3BP1 exhibited characteristics typical of SGs, a distinct form of LLPS. Furthermore, we demonstrated that the knockdown of LINC00599 using siRNA disrupted SG formation and inhibited the proliferation of PASMCs under hypoxic conditions. Our findings indicate that LINC00599 plays a role in hypoxia-induced proliferation of PASMCs through the formation of LLPS with G3BP1. To the best of our knowledge, this study is the first to demonstrate that lncRNAs mediate PASMC proliferation via LLPS.

Myosin heavy chain 9 (MYH9), a member of the myosin family of proteins, is a skeletal protein that plays a critical role in many human diseases by regulating cell proliferation^38–40^. However, the role of MYH9 in PH is still unknown. Using liquid chromatography-tandem mass spectrometry (LC-MS/MS), we selected MYH9, the top protein on the list, for further study as the downstream protein. Through bioinformatics analysis, we predicted that the MYH9 protein possesses intrinsically disordered regions and has the potential to form LLPS. Subsequently, we proved that MYH9 bound to LINC00599 and was involved in the LLPS structure. Additionally, LINC00599 positively regulated the expression of MYH9. Knockdown of MYH9 with siRNA also disrupted the formation of LLPS and restrained the proliferation of PASMCs under hypoxic conditions; this effect was reversed when co-transfected with the wild LINC00599 plasmid. These data reveal that LINC00599 regulates the prolieration of PASMCs through LLPS with MYH9 under hypoxic conditions.

The mechanisms by which lincRNAs mediate the proliferation of PASMCs are not fully elucidated. There is a research gap in the epigenetic modification of lincRNAs and their effects on the proliferation of PASMCs. N6-methyladenosine (m6A) is the most abundant chemical modification of RNA found in mRNAs and noncoding RNAs^41^. Nevertheless, the role of m6A modification in regulating LLPS formation remains unknown. Through bioinformatics analysis, we predicted that the binding sites of LINC00599 and G3BP1 overlap with the m6A sites of LINC00599. The mutation of these m6A sites disrupted the binding of the two molecules and affected the LLPS structure. Moreover, the overexpression of LINC00599-Me-mut could not rescue the proliferation of PASMCs inhibited by LINC00599 siRNA. Overexpression of LINC00599 with the m6A site mutations failed to reverse the PH improvement induced by LINC00599 shRNA in vivo. These data reveal that the m6A modification of LINC00599 regulates the formation of LLPS and the hypoxia-induced proliferation of PASMCs.

Super-enhancers (SEs) are large enhancer clusters, and the activation of specific SEs is associated with transcriptional dysregulation of ncRNAs in the course of diseases^42,43^. We predicted that the expression of LINC00599 was associated with SE-2082886 and filtered out the key transcription factor ZNF263, which mediates the regulation of SE on the LINC00599 gene promoter through bioinformatics analysis.

Subsequently, we confirmed the regulation of SE-2082886 on the gene promoter of LINC00599 via ZNF263. Moreover, we illustrated the enhancer 2 and promoter (nucleotides −1250 to −1500) are the key regions regulating the expression of LINC00599. Our results revealed the key molecules required for SE-driven LINC00599 expression, as well as the binding domains and sites between SE-2082886 and the LINC00599 gene. Therefore, our results provide more comprehensive evidence of an upstream mechanism driving the expression of the LINC00599 gene.

In summary, we demonstrated, for the first time, a novel role of LINC00599 in the pathogenesis of PH. The function of LINC00599 in the development of PH is likely mediated through liquid-liquid phase separation with G3BP1 and MYH9 in the cytoplasm of PASMCs. Additionally, the upregulation of LINC00599 may be due to SE-2082886 mediated by ZNF263. These findings indicate that LINC00599 may be an important target for therapeutic interventions in hypoxia-induced PH.

## Materials and Methods

### Animals

To eliminate the possible effects of estrogen on pulmonary arterial hypertension, this study exclusively involved 6-week-old male C57BL/6J mice, with a body weight ranging from 20 to 25 grams and in good health, sourced from the Experimental Animal Department of Harbin Medical University. Before the commencement of the experiments, the mice were acclimated in a specialized animal facility and provided with ad libitum access to food and water for a period of 10 days. The animal care and utilization protocols strictly adhered to the “Guide for the Care and Use of Laboratory Animals” as published by the National Institutes of Health (NIH Publication No. 85-23, revised in 1996). All animal procedures were approved by the Medical Animal Ethics Committee of Harbin Medical University - Daqing Campus (Daqing, China) (NO: HMUDQ20241029002).

### Rodent PH Models

The mice were randomly grouped. The Sugen5416-induced hypoxic pulmonary hypertension model (SuHx) in C57BL/6 mice was established by exposing them to a normobaric hypoxic environment (FiO_2_ 0.10) for 21 days, along with intraperitoneal injections of Sugen5416 (20mg/kg) on days 1, 7, and 14. Following the induction of the hypoxia-induced pulmonary hypertension model, the mice were maintained under normoxic conditions (FiO_2_ 0.21) for an additional 28 days.On the first day of the normoxic environment, the mice received lentivirus (LVV) viral particles carrying LINC00599 shRNA (sh-LINC00599), a short hairpin RNA targeting LINC00599, or scrambled shRNA (NC) sequences via nasal instillation. After 14 days, the mice were administered another set of LVV viral particles via nasal instillation, this time carrying a mutated version of LINC00599 RNA with the m6A site mutated (LINC00599-Me-mut). The viral particles were constructed by GENECHEM (Shanghai, China), and a vector suspension containing 10^10^∼10^11^ genome equivalents in 20-30 μl of Hanks’ Balanced Salt Solution was prepared for administration of LVV viral particles. The study was approved by the Harbin Medical University (Daqing) Ethical Committee. All animals were housed in a temperature-controlled environment with a 12:12-hour light-dark cycle. Surgical procedures were conducted under tribromoethanol anesthesia to ensure minimal pain.

LINC00599 shRNA: 5’ - GCAACCAGGATCCTTTAAAGG - 3’

Scramble shRNA: 5’ - CCGATCTGACATGACTGCG - 3’

LINC00599-Me-mut:

CGCCCTCCGCAGACCTCCGCGCAGCGGCCGCGGGCGCGAGGGGAGGGGTCTGGAGCTCCCTCCGGCTGCCTGTCCCGCACCGGAGCCCGTGGGGTGGGGAGGTGTGC AGCCTGTGACAGACAGGGGCTTAGAGATGCAA**T**CAG**T**CTCAGGGAGAGAAACAGA AGCTGATTCTGTGACAGAAGCAGATCTGTGCAGCACAGATGCGGTGTGCGTGGGGAGGGGGTCGCCTGGGAGCGCATTGCGGAGTGCTTGTGTGTGCAGATTTTTCTCTGGGCTCAGGACTCATTGTATGTGGGTCAACACCTTCCTCCGTGACTGTGTTTTTGTTCTGAGCTGAGTTTTTTGGTTTGCCCTTAAAAAAATAATAATTTGGCATCCAGAGACTGG CAGACTGCCTCAGGGCCTGG**T**CTGCGGATATATTGTGTTCTGCTTGAGGTTTGGGG AGGAGGGCAGGCGGTAGGAAGGGAGAGGGGGAGCTGTTTGTCACACTTTGCTGT AGAGCTGAGAGCACCTGACAAGCTTAAGGAAGTCGTTGGGCTATGTGGACAAGAAGGAGCCAGCTCCCAGCGGGTTCACAAGCTCTATCGGAGTTGAAAGCGTGGTCATGGCTCTAAGGAGCACCTCACGCCCTCCCTGTAGCTGTTATTGCAGTTTCAGGCAGAGATCCAGGAGCTGCAGAGGAAGGGAGAGGCACAATAACCTACATGGACCCAAGGGAGACATGTGTTCCTTTAAAAATGTGAACAGAAGGAAAAACAGAATGTGTGCAACTGGGGGTCTGAGGAAAGACTGTTTTGAAAGAGGCTGTCAGGGAATGGAAAGGGTTAAGCTTTTCATCCTGAAGAACCTGCTTCCCAAATCAGGCCTTCCTCCCATCACTAGACCCTGAGCAGCAGCTGGTCCTAGAGACCCCCCTCGTACTGCGCTGCCACAGTCTCATCCCATTTCCAGCTCTGTATTAACCACCAAGCTGCAGCGGATGGGGCAAGACCTAAGCTCATTAATTTCCAGGTGGAGGAAATGGAGGAGGGGCAGGTCCCTGCAGTAGAGGGAGGAGCAGAGAAAGGAGGCCAAGAGCTGGATCCTTCTGCCCAGGAAGCCTGCTGCATCCCTTCCCCCGAGCATGGCAGAGGCCTGGCTTTGCAAGGCCAAGGCCATAAGGGATGCTTAGGAGATTAATTTGATTCCTGACACAATAATCAAGCCCTAAGAGTCTCCACTGAAGCTTACTGAGGACTTCTTTCCTCTCCAAAGCCTCAGTCTAGCCTGCTAAATAAATTAGTATTCAGTGATGCCTTGGATCAGGGCCCCTCCCCGGCCTCAGTTTCCCCAAATATTTATTAAGTACCTACTGTGTGCAATCCTTGTGTAATTATTACCTCTTAGGCTCTTCATTTGCCCTCCTAAAGCAGTGTTTAGAGTCAGGCAGAGGTTAAGTGTGTTGCACCTCACCTAGATACTTCCAGAACCTTCTCTGGGTCTGCAGAATGTGGCACAACCTGCTTGCCCCCGCAGAGAGAAAGCTGCAGTGCACATCCTGCAGACTGCAGGTGCTGGGCTGCCTCTGGAGTCCCAGAAGGCAAGCTTGGCTGCAGGACAGAAAGGGAGAACAGCTTCTCTCACCCCTGAGCCTTCACAAGCCCTTGTCTATTTGCCGTTGCCTTCAAAATATACCTCCCCCGAAACCAGTAGCTTTCTGAGTCCTGGTGTCCCCTCCGCCCTTTCTGGACAGGTTTGGGAAGAAGAAAGCAGTCAGTGCTGGGCCTTATTGGGGTGTGAAG CGCCTTGCTCTGCCCCTTCTGCTCACTGTGAAGGCCGCTGGATGCTTCTCTTAGGCATGGTTTAAGCCTCCGATTACTAAACCCCTTGCCCCACAAACGTCCACATTGACGAGCCTCTTTTTAGTAACTGCTTCCCCGTAATTCCTTCAGAGGTTGCTGTACCCTTCGCTGATGTGCTGCCCTCCTGTAAAACCTCCAGATGCCTTCCCACGTAATGCCCCTTTCAGATGCTTTAAGCTGAGAGCTTAAACCACAGGTACCATGGCTGACGCCTGCCAGGTTTCTGCTGCAGATAATCTATGATGGGAGGGGCATATTTTTTACTTCATTACTTATGTAAACTCTTGTTCCAGAAAGCTTTAATGTGTGTGGGAGTGTTCTGGGTCTATTAGGTCTGTGCGCATGGGTGTGGGCATTTGCCTGTGTCCACCGGGTGGGTCTCATTATGAAATGTATGTTTATGTAGGGCTTTAATGGCTGAAAATGGCAAAGAGATGAATAGACCACTTGGCCCCATGTGTAATTGCCAGGCCCCTTCTGTGCTCAAATGAGGTGTCCGAGTGAAGGTCAGCCCTTCCCTTCTGTATTTGGGGCCTATTTATGCCACCAGTAATTTTATAAGAAATCTGAATAGTTCTCCCCTTTGAGTGCATTTAACTCTTTAGTATCTTCTCTCTTACCTATTTGAGCCCCTCTAGCTACAGTCTGGCTTAAATGAAAGGGGAATTATATGCTTAAGAAAAAGTAGGACACGGTTGAGGCAGTTTGCTGACTGAATACGCGAAGAAGGACCTGATGGGCTCATATGCACCACTGCCATCACAGTCCCCATCGTGATGCAAGCTTATATGATTCTTGAGAAACAGCCCAGATCTCCTCTTAAGCGGAAAAGAGATTGACCTTCTAGCAGAGGCAAAGAATGGAGTCTTCTGGGGGCAGCTGCAAAGCGTTCTCCCTAGGACAGATGGAGCCTCCCTTTCCTCATCTACTCTGTGGGTGGTTTCAGGGCCCACGAGTCAACATGAGGAGTTGTGCTGGTGGTATGTGTGTTGGAGGCTGGGCTGGCTGATTCACAGTGACGAGGATGTCAATAATAACAAGAATGAGAATGATGGTACCTAATAAAGACTTTTTTTCCCAA

### Echocardiography

Mice were anesthetized with amobarbital (60 mg/kg, i.p.) and evaluated using a Vevo2100 ultrasound system for cardiac structure and function. Measurements included the pulmonary arterial velocity time integral, preejection time, and ejection time. The operator did not know how the mice were grouped.

### Hemodynamics Measurement and Tissue Preparation

The FA-404 recorder was used with a 1.2 French catheter (Scisense, Inc) to measure right ventricular pressure. Following this, the heart was dissected and weighed to calculate the right ventricular hypertrophy index (RV/LV+S). Hemodynamics were measured by professionals who did not know the grouping.

### Culture of PASMCs

In this study, the human pulmonary artery smooth muscle cells (HPASMCs) were procured from Hunan Fenghui Biotechnology Company, (Changsha, China). These cells were cultured in a specific smooth muscle cell medium (SMCM) and passaged for subsequent experiments. The culture conditions were maintained at a temperature of 37°C and a carbon dioxide concentration of 5% in a humidified atmosphere. To mimic hypoxic conditions, the cells were further cultured in a tri-gas incubator manufactured by Heal Force (Shanghai, China), with a gas mixture composed of 3% oxygen, 5% carbon dioxide, and 92% nitrogen.

### Cell Transfection

GenePharma Company (Shanghai, China) was responsible for designing and synthesizing the siRNA and its negative control (NC). Additionally, Genechem Company (Shanghai, China) designed and synthesized the overexpression plasmids and the empty vector. At the start of the experiment, cells were seeded in culture dishes and cultured until they reached 60∼70% confluence. The transfection reagent, siRNA, NC (negative control), and plasmids were then diluted in DMEM. Subsequently, the diluted reagent was combined with the diluted siRNA, NC, and plasmids to create transfection complexes, which was then added to the cell culture medium. Six hours post-transfection, the cells were placed in a hypoxia chamber (37°C, 3% O_2_, 5% CO_2_, tri-gas incubator, Shanghai, China) to simulate a hypoxic environment. After an additional 48 hours of culture, experimental procedures were carried out. Sequences for siRNA were as follows:

LINC00599 siRNA-1: 5′ - GGAUGCGGGAGAACAAAGATT - 3′

LINC00599 siRNA-2: 5′ - UCACAGCGGACCUUGAUUUTT - 3′

MYH9 siRNA-1: 5’ - GCAAGCUGCCGAUAAGUAUTT - 3’

MYH9 siRNA-2: 5’ - GCAAAUUCAUUCGCAUCAATT - 3’

G3BP1 siRNA: 5’ - GGGCUUCUCUCUAACAACATT - 3’

ZNF263 siRNA: 5′ - GGAUAUGCAGAGAGAGCUUTT - 3′

### Western Blot

RIPA lysis buffer (Beyotime, China) was used to extract tissue or cell proteins, and 10%∼12% polyacrylamide gels were prepared for protein separation. After transferring the proteins to the nitrocellulose membrane, it was blocked in a buffer (20 mM Tris, pH 7.6, 150 mM NaCl and 0.1% Tween 20) containing 5% nonfat dry milk. The membranes were incubated overnight at 4 °C with primary antibodies and then reacted with the appropriate horseradish peroxidase-conjugated secondary antibodies. The specific bands were visualized through enhanced chemiluminescence (M2301, HaiGene, China).

### Reverse transcription and Real-time PCR(qRT-PCR)

Total RNA was extracted from cells or tissues using Trizol reagent (Invitrogen, USA). RNAs in the cytoplasm and nucleus were isolated using a PARISTM kit (Thermo Fisher Scientific, CA) following the manufacturer’s instructions. Complement DNA was generated using the PrimeScript™ RT reagent Kit (TaKaRa, Japan). Real-time PCR was performed on a LightCycler 480 II real-time PCR system (Roche, Germany) with TB Green® Premix Ex Taq™ (TaKaRa, Japan) to measure RNA expression. β-actin RNA was used as an internal control for total RNA quantification. 18S RNA was used as an internal control for cytoplasm RNA quantification, while U6 served as an internal control for nuclear RNA quantification. The threshold cycle (Ct) was determined, and relative RNA levels were calculated based on the Ct values. Data analysis was conducted using the 2^−ΔΔCT^ method.The primers were as follows:

LINC00599 forward: 5′ - CAACACCTTCCTCCGTGACTGTG -3′

LINC00599 reverse: 5′ - GCTGGCTCCTTCTTGTCCACATA -3′

β-actin: forward: 5′ - GCACTCTTCCAGCCTTCCTT - 3′

β-actin reverse: 5′ - TGTGTTGGCGTACAGGTCTT - 3′

18S RNA: forward: 5′ - CGGACACGGACAGGATTGACAG - 3′

18S RNA reverse: 5′ - AATCGCTCCACCAACTAAGAACGG - 3′

U6:forward: 5′ - CGCTTCGGCAGCACATATAC - 3′

U6 reverse: 5′ - TTCACGAATTTGCGTGTCATC - 3′

MYH9 forward: 5′ - ATCCTGGAGGACCAGAACTGCA - 3′

MYH9 reverse: 5′ - GGCGAGGCTCTTAGATTTCTCC - 3′

### RNA Immunoprecipitation (RIP)

Binding between RNA and protein was analyzed using RNA Immunoprecipitation kit (Bes5101, Bersinbio, Guanzhou, China) according to the manufacturer’s instructions. In summary, the process involved using antibodies specific to the target protein to precipitate the corresponding RNA-protein complexes. Reverse transcription was performed or a cDNA library was constructed. Ultimately, qRT-PCR was utilized to analyze the types and quantities of RNAs associated with these complexes.

### Chromatin immunoprecipitation (ChIP) assay

ChIP Assay Kit (P2078, Beyotime, China) was utilized, following the manufacturer’s protocol. The process began with cross-linking cells to formaldehyde, followed by the addition of lysis buffer and sonication to shear the genomic DNA. Anti-H3K27ac antibody (A2363, ABclonal, Boston, USA), anti-H3K4me3 antibody, or anti-ZNF263 antibody were then added and incubated with protein A+G Agarose/Salmon Sperm DNA. An anti-IgG antibody was used as a negative control to capture the target gene sequences for PCR amplification. The cross-linked DNA released from protein-DNA complexes was purified using a DNA extraction kit (D0033, Beyotime, China), and the eluted DNA was analyzed by qRT-PCR. The specific primers used for ChIP-qPCR were:

LINC00599 promoter 1 forward: 5′ - CGCTGCTCTCCTTGTCCTT - 3′

LINC00599 promoter 1 reverse: 5′ - CTTCGCCCAGGAAACTGAAG - 3′

LINC00599 promoter 2 forward: 5′ - GAGTGGAGATGGAGGTGAGT - 3′

LINC00599 promoter 2 reverse: 5′ - CCTTCTCCTCCTCAGTGCTC -3′

LINC00599 promoter 3 forward: 5′ - CGCTGCTCTCCTTGTCCTT - 3′

LINC00599 promoter 3 reverse: 5′ - CTTCGCCCAGGAAACTGAAG - 3′

Enhancer 1 forward: 5′ - ACACTACGGAAAAGGGAGCA - 3′

Enhancer 1 reverse: 5′ - AGGGTTTTTAGCTTCAGGACAAT - 3′

Enhancer 2 forward: 5′ - AAATGGCCTCGAAGCGAGAA - 3′

Enhancer 2 reverse: 5′ - AGGGCAACTAGAGATGGGGA - 3′

Enhancer 3 forward: 5′ - AGCCCTAAAACGCGCCG - 3′

Enhancer 3 reverse: 5′ - GCTTAGGCAGACGGAGTGG - 3′

Enhancer 4 forward: 5′ - ACGCTACCAAGACCTGAGGG - 3′

Enhancer 4 reverse: 5′ - ACCCTCCCACTGTAATTTTGAGA - 3′

Enhancer 5 forward: 5′ - CATTTCGGGCCCCGTCG - 3′

Enhancer 5 reverse: 5′ - GAAAAGGGAAACCAGCGCCT - 3′

Enhancer 6 forward: 5′ - GATCCACTTCAGTACCTGCG - 3′

Enhancer 6 reverse: 5′ - CAAGCGCTTTCGCGGC - 3′

### Co-immunoprecipitation (Co-IP) assay

After setting up the cell model, cells were washed with cold PBS, treated with 1 ml of lysis buffer (Beyotime, P0013B) plus PMSF (1:100), and collected in a 1.5 ml tube. The tube was placed on ice and shaken gently for 30 minutes. After centrifugation at 4°C and 15,000 rpm for 30 minutes, 100 μl of the supernatant was set aside as a blank control. The rest was mixed with 40 μl of washed protein A+G magnetic beads (Beyotime, P2108) and incubated at 4°C for 1 hour with slow shaking. Following magnetic separation, 400 μl was divided, with 3 μl of target or IgG antibody added to each, and incubated at 4°C overnight. An additional 20 μl of magnetic beads was added, and the mixture was incubated at room temperature for 1 hour. The beads were washed with 500 μl of 1X TBS (pH 7.5). SDS-PAGE loading buffer was then added, heated at 95°C for 5 minutes to elute proteins, and the supernatant was used for western blot analysis.

### RNA Pull-Down

Binding between RNA and protein was analyzed using RNA pull-down Kit (Bes5102, Bersinbio, Guanzhou, China) according to the manufacturer’s instructions. A biotin-labeled LINC00599 probe was generated using in vitro transcription. This probe was then incubated with a cell total protein extract to form RNA-protein complexes. These complexes can bind to streptavidin-labeled magnetic beads, allowing them to be separated from other components in the incubation mixture. After elution of the complex, mass spectrometry (Beijing Bio-Tech Pack Technology Company Ltd., China) was employed to identify proteins that bind to LINC00599. Furthermore, western blot analysis was conducted to determine whether G3BP1 or MYH9 interact with LINC00599.

### Flow Cytometry

After establishing the cell model, the cell culture medium was collected and the cells were digested with trypsin. The previously collected medium was then added to disperse and resuspend the cells. The cells were centrifuged at 1000g for 5 minutes to form a pellet. The majority of the supernatant was removed, retaining about 50 microliters, and the cells were resuspended in cold PBS and centrifuged again to wash. Subsequently, the cells were fixed with cold 70% ethanol and stored at 4°C. Prior to staining, a suitable quantity of propidium iodide staining solution was prepared following the guidelines of the Cell Cycle and Apoptosis Detection Kit (C1052, Beyotime, China), and 0.5 ml of this staining solution was added to each tube. The tubes were incubated at 37°C in the dark for 30 minutes. Flow cytometry analysis was conducted to determine the proportions of cells in the G0/G1 phase, S phase, and G2/M phase after the staining process was complete.

### Cell Viability and Proliferation Assays

Utilizing the Cell Counting Kit-8 (C0037, Beyotime, China) for evaluating cell viability and proliferation, a density of 2,000 to 3,000 cells per well was plated into a 96-well plate, ensuring an even distribution for accurate assessment. After the completion of the treatment, 10 μl of CCK-8 reagent was added to each well, and the plate was incubated at 37°C for 3 hours. The absorbance was then measured at a wavelength of 450 nm.

### Immunofluorescence

Prepared lung tissue sections or processed cells were fixed in 4% paraformaldehyde, permeabilized using 0.3% Triton X-100, and blocked with 5% bovine serum albumin at 37°C for 1 hour. Subsequently, the tissues or cells were incubated with anti-Ki-67 antibody (1:100) at 4°C overnight. After washing three times with PBS, fluorescently labeled secondary antibody was used to incubate for 2 h at 37°C. After washing three times with PBS, DAPI was incubated at room temperature for 15 minutes. Confocal laser scanning microscope (CLSM) was used to acquire images.

### 5-Acetylidene-2’-Deoxyuracil Nucleoside (EdU) Assay

The proliferation of cells was assessed using the BeyoClick™ EdU-488 Cell Proliferation Assay Kit (Beyotime, C0071S) following the manufacturer’s protocol. Cells were plated in a 24-well plate, and the EdU reagent was added to achieve a final concentration of 10 μM and incubated at 37°C for 2 hours to label the cells. Cells were then fixed in a 4% paraformaldehyde solution for 15 minutes and permeabilized with 0.3% Triton X-100 for 15 minutes. Subsequently, the cells were incubated with the Click reaction mixture at room temperature in the dark for 30 minutes. Cell nuclei were counterstained with 1× Hoechst 33342 reagent. The staining results were observed under a fluorescence microscope.

### Fluorescence in Situ Hybridization (FISH)

A cell or paraffin section FISH kit (GenePharma, Shanghai,China) was used to perform quantitative and localization analysis of RNA, following the manufacturer’s instructions. Essentially, the process involves using a known fluorescent dye to label single-stranded nucleic acids as probes. These probes bind specifically to the target RNA based on the principle of base complementary pairing. After denaturation, annealing, and hybridization, a hybrid complex of the target RNA and the nucleic acid probe is formed. This complex can then be observed under a fluorescence microscope.To detect the co-localization of RNA and proteins, the antibodies for the protein of interest and the corresponding fluorescently labeled secondary antibodies need to be incubated before nuclear staining.

### Fluorescence Recovery After Photobleaching (FRAP)

293T cells were cultured in glass-bottom dishes (Nest, 801002) and transfected with a G3BP1 fluorescent plasmid. For imaging, a confocal microscope was used to visualize the cells. A 405-nm laser was applied to bleach the fluorescence signal, allowing for the continuous capture of images and the measurement and recording of fluorescence intensity over time.

### Hematoxylin-Eosin Staining (HE)

The paraffin sections were deparaffinized by dissolving the paraffin, quickly immersing the baked sections in xylene for two 5-minute dewaxing treatments. Then, a gradient hydration process was carried out, placing the sections in 100%, 90%, 80%, and 70% ethanol for 5 minutes each, and finally in distilled water for 5 minutes. After drying, hematoxylin staining solution was added for 5 minutes, rinsed with running water, and the degree of nuclear staining was observed under a microscope. A 1% hydrochloric acid alcohol solution was used for 1 to 3 seconds, rinsed, and then placed in tap water to blue for 5 to 10 minutes, adjusting the staining and differentiation times based on microscopic observation. For counterstaining, 0.5% eosin solution was added dropwise, stained for 10 to 15 seconds, and observed under a microscope after rinsing with distilled water. Subsequently, gradient dehydration was performed with 80%, 90%, 95%, and 100% ethanol. The sections were cleared with xylene twice for 5 minutes each and air-dried at room temperature. Finally, a drop of neutral balsam was added to the tissue, covered with a cover slip, and then observed and photographed under a microscope.

### Statistical Analysis

Statistical analysis was performed with GraphPad Prism 8 software. Data were checked for normal distribution and equal variance (F test) before statistics. Students t test (unpaired) was used for 2-group analysis with equal variance, and Welch correction test was used for 2-group analysis with unequal variance. One-way ANOVA with Tukey post hoc test was used to compare multiple groups with equal variance, and Brown Forsythe and Welch ANOVA with Tamhane T2 post hoc test was used to compare multiple groups with unequal variance. Nonparametric analyses, including the Mann-Whitney U test for 2 groups or Kruskal-Wall is test followed by Dunn post-test for >2 groups, were performed for non-normally distributed data. Data are presented as the means±SEM, and P<0.05 was considered statistically significant.

## Nonstandard Abbreviations and Acronyms

CDK: cyclin-dependent kinase
CyclinA: G2/M-specific cyclin-A2
CyclinE: G1/S-specific cyclin-E1
DAPI: 4’,6-diamidino-2-phenylindole
EdU: 5-Acetylidene-2’-Deoxyuracil Nucleoside
G3BP1: Ras GTPase-activating protein-binding protein 1
H3K27ac: Histone H3 Lysine 27 acetylation
H3K4me3: histone H3 lysine 4 trimethylation
hPASMC: human pulmonary arterial smooth muscle cell
IDR: intrinsically disordered region
LINC00599-Me-mut: lentivirus carrying LINC00599 with m6A sites mutation.
IncRNA: long non-coding RNA
LVV: lentiviral vectors
mPAEC: mouse pulmonary artery endothelial cell
mPASMC: mouse pulmonary arterial smooth muscle cell
MYH9: myosin heavy chain 9
PAH: pulmonary arterial hypertension
PAT: pulmonary artery acceleration time
PAVTI: pulmonary artery velocity time integral
PCNA: proliferating cell nuclear antigen
PH: pulmonary hypertension
RVH: right ventricular (RV)/ left ventricular (LV )+ S weight ratios
RVSP: right ventricular systolic pressure
sh-LINC00599: shRNA-mediated silencing of LINC00599
ZNF263: zinc finger protein 263
α-SMA: α-smooth muscle actin

## Acknowledgements

We would like to thank Associate Professor Baoshan Zhao for providing guidance for the pathology platform of Harbin Medical University (Daqing).

## Author Contributions

H.Y. and D.Z. designed research; Y.W., L.Y., S.Z., A.L., C.L., Z.Z., X.Z., Y.P., Z.B, and H.Z. performed research; H.Y. contributed new reagents/analytic tools and funding support; Y.W., L.Y., S.Z. and A.L. analyzed data; and H.Y., Y.W. and L.Y. wrote the paper.

## Data and materials availability

All data needed to evaluate the conclusions in the paper are present in the paper and/or theSupplementary Materials. The mechanic diagram is made by Figdraw.

## Perspectives

- Elevated levels of LINC00599 in murine pulmonary hypertension (PH) models indicate its potential utility as a biomarker.
- The silencing of LINC00599 attenuates experimental PH by disrupting the formation of stress granules, thereby underscoring its promise as a therapeutic target for PH.
- RNA m6A modification has been associated with various cardiovascular diseases; however, its involvement in liquid-liquid phase separation (LLPS) has not been thoroughly investigated. This study demonstrates that LINC00599 regulates LLPS through m6A modification, influencing the proliferation of pulmonary artery smooth muscle cells (PASMCs). Moreover, overexpression of LINC00599 with mutated m6A sites does not alter the enhanced effects observed with LINC00599 shRNA in SuHx mice.
- Deletion of LINC00599 is sufficient to reverse the progression of PH; however, the upstream regulatory mechanisms remain poorly understood. Our research has identified the super enhancer (2082886) as a critical upstream regulator within the genome that facilitates the expression of LINC00599 through the transcription factor ZNF263.

## Supplementary Materials

**Figure S1.**
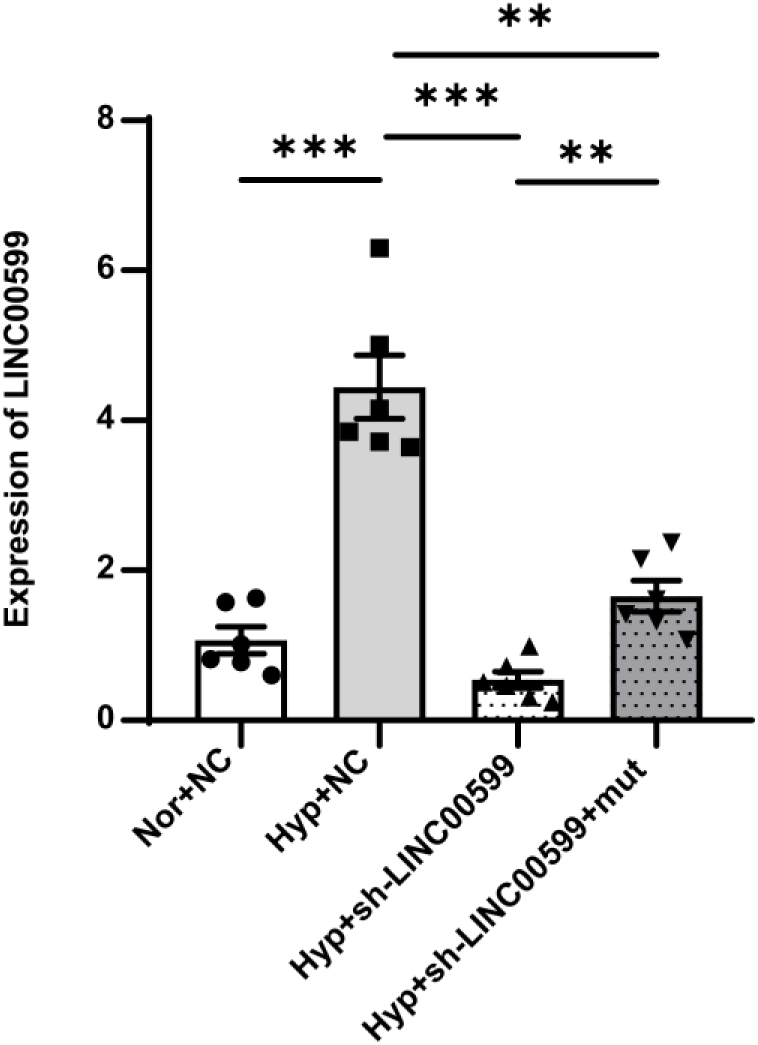
The effects of LINC00599 silencing via shRNA and overexpression through plasmid introduction. RT-qPCR showing the levels of LINC00599 in mice (n=6). Data were expressed as mean ±SEM. *\*\*, P<0.01; ***, P<0.001.* Nor, normoxia; Hyp, hypoxia; NC, negative control; sh-LINC00599, lentivirus carrying LINC00599 shRNA; mut , lentivirus carrying LINC00599 with m6A sites mutation.

**Figure S2.**
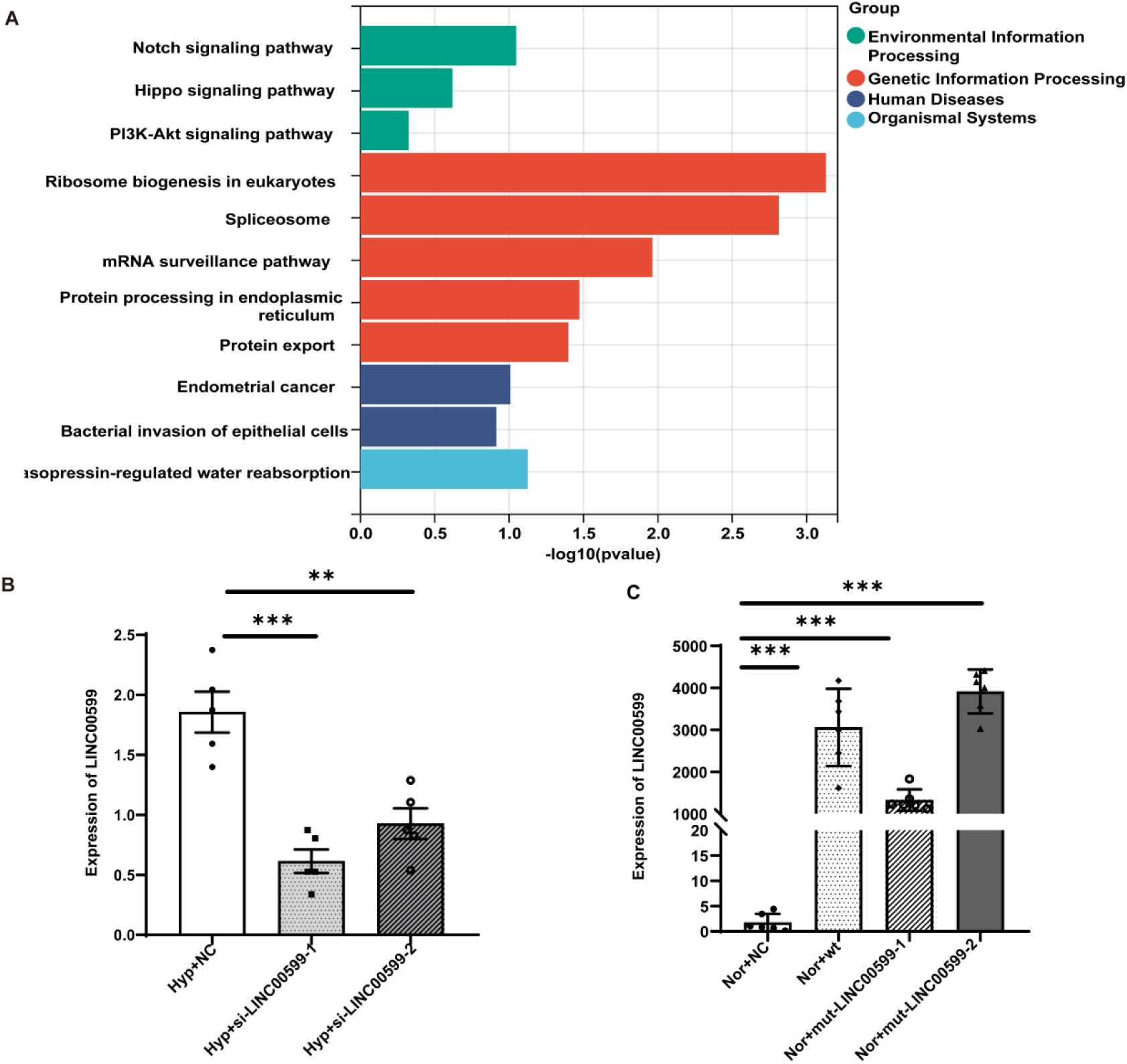
The analysis of KEGG pathways and the impact of LINC00599 siRNAs and overexpression plasmids. (A) KEGG enrichment analysis of LINC00599. (B) qRT-PCR assay showing the silencing effect of si-LINC00599 in hPASMCs (n=5). (C) qRT-qPCR showing the overexpression effect of LINC00599 plasmids in hPASMCs (n=6). Data were expressed as the mean ± SEM. *\*\*, P<0.01; ***, P<0.001.* Nor: normoxia; Hyp: hypoxia; NC: negative control; wt: overexpressing wild type LINC00599 plasmid; si-LINC00599: LINC00599 siRNA; mut-LINC00599: lentivirus carrying LINC00599 with m6A sites mutation.

**Figure S3.**
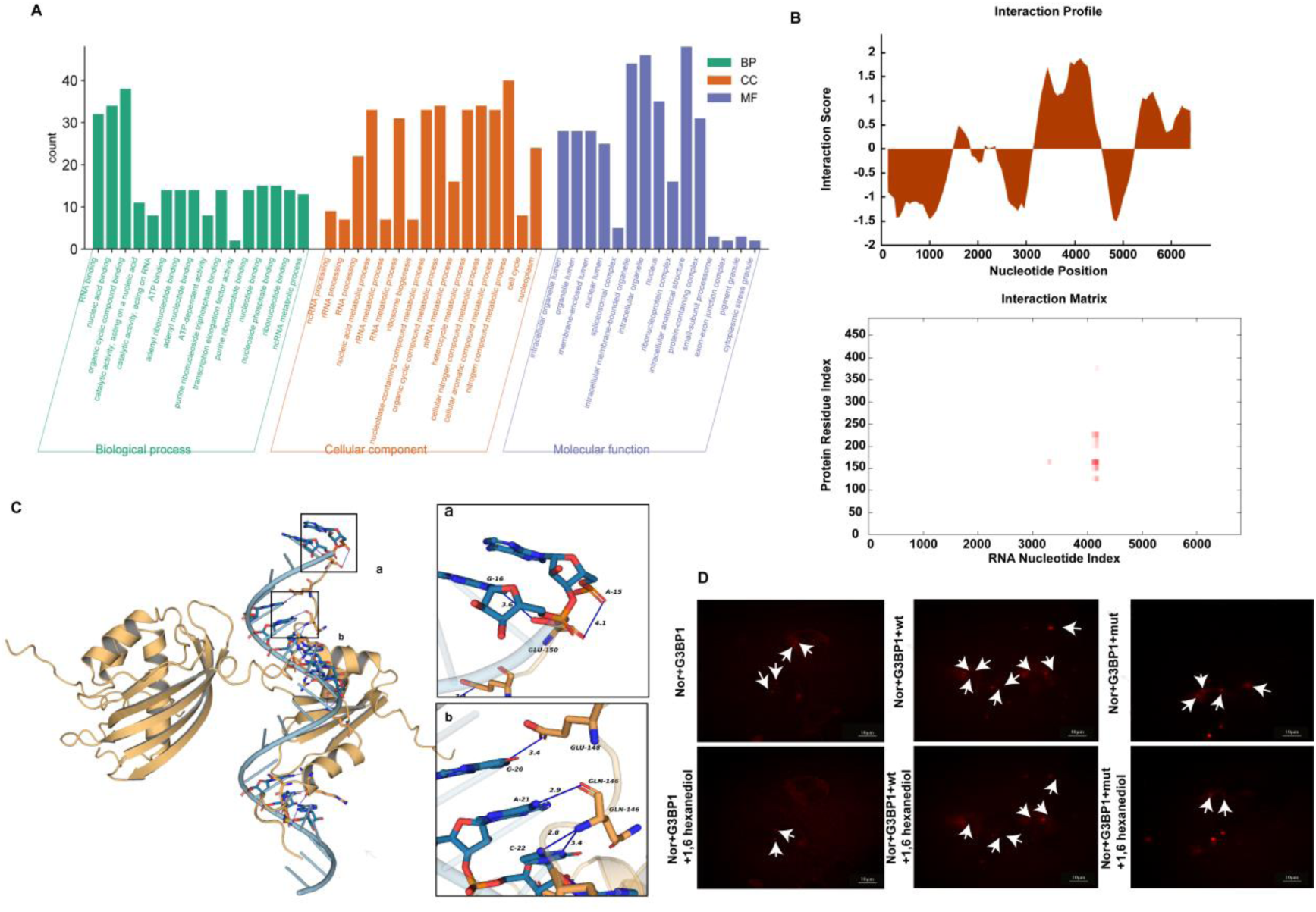
The analysis of GO pathways and the binding interaction between LINC00599 and G3BP1 in hPASMCs. (A) The Gene Ontology analysis elucidating the biological functions of LINC00599. (B) Bioinformatic predictions identifying proteins that potentially interact with and bind to LINC00599. (C) Molecular docking studies analyzing the binding sites between LINC00599 and G3BP1. (D) FISH and immunofluorescence assays demonstrating the interaction between LINC00599 and G3BP1 in 293T cells following treatment with 1,6-hexanediol. Nor: normoxia; wt: overexpressing wild type LINC00599 plasmid; mut-LINC00599: lentivirus carrying LINC00599 with m6A sites mutation.

**Figure S4.**
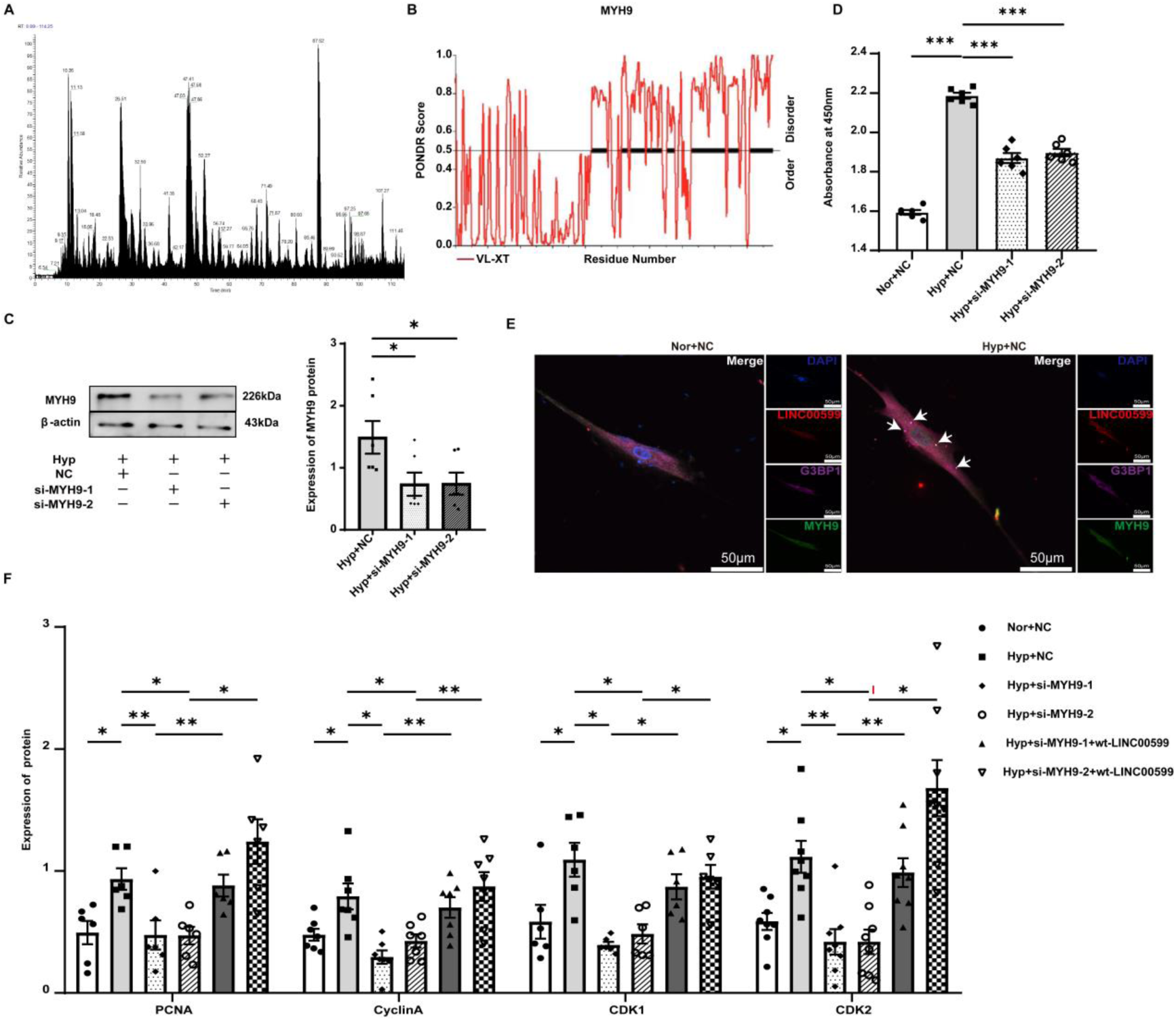
MYH9 interacts with LINC00599 and enhances the hypoxia-induced proliferation of hPASMCs as a downstream effector of LINC00599. (A) LC-MS/MS analysis identifying proteins that interact with LINC00599. (B) Bioinformatic predictions revealing intrinsically disordered regions within the MYH9 protein. (C) Western blot analysis demonstrating the silencing efficiency of MYH9 siRNA (n=6). (D) CCK8 assay assessing the impact of MYH9 siRNA on hPASMC viability (n=6). (E) FISH and immunofluorescence assays illustrating the co-localization of LINC00599 and G3BP1 with MYH9 (n=3). Scale bar: 50μm. (F) Western blot analysis quantifyings the expression levels of proliferation-related proteins (n=6). Data are expressed as the mean ± SEM. *\*, P<0.05; **, P<0.01; ***, P<0.001.* Nor: normoxia; Hyp: hypoxia; NC: negative control; wt-LINC00599: overexpressing wild type LINC00599 plasmid; si-LINC00599: LINC00599 siRNA; si-MYH9: MYH9 siRNA.

**Figure S5.**
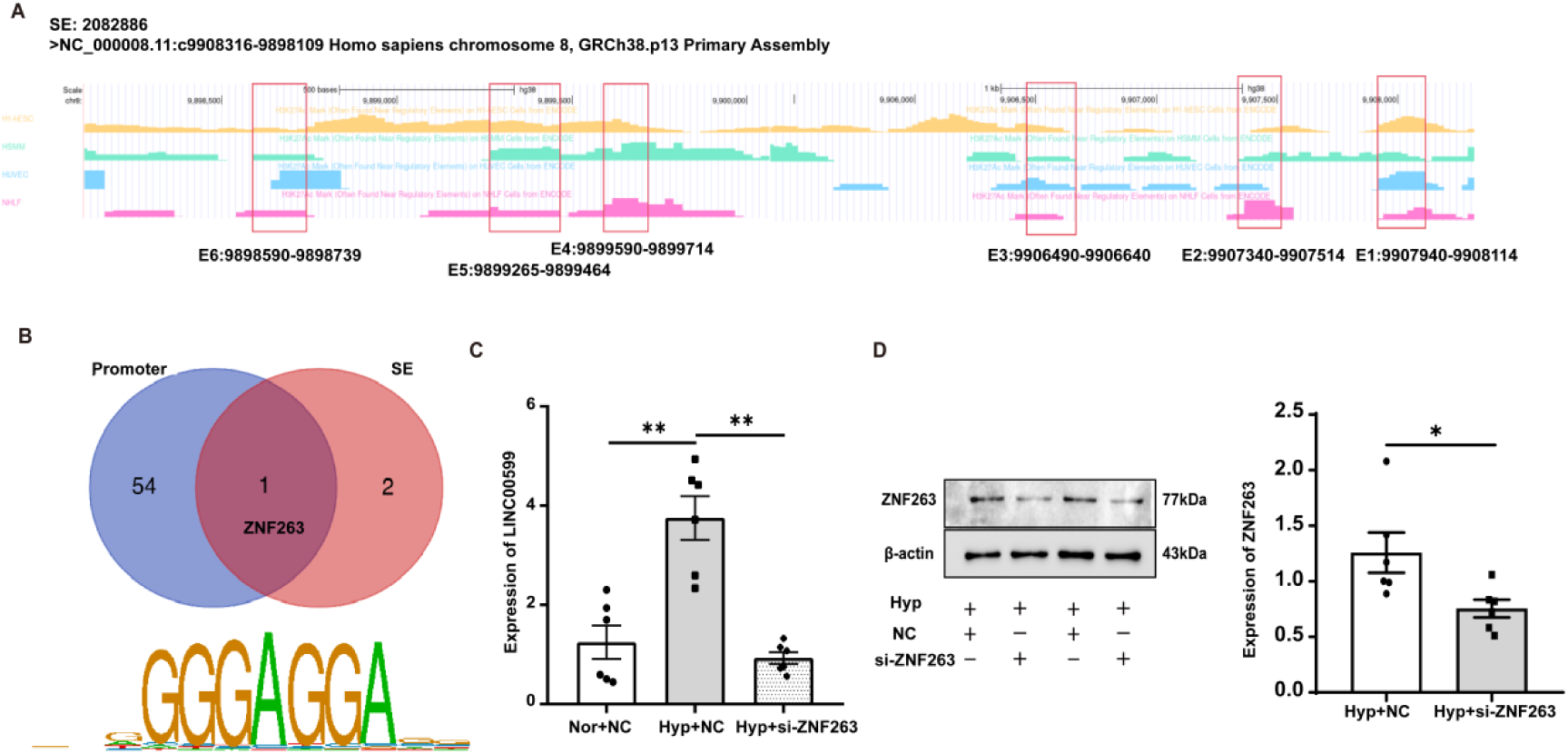
The composition of SE-2082886, the analysis of ZNF263, and the enrichment of the H3K27ac region within the LINC00599 promoter. (A) The composition of the super enhancer regulating the expression of LINC00599 by the UCSC database. (B) The screening process of transcription factor using bioinformatics. (C) qRT-PCR showing the silencing effect of ZNF263 siRNA (n=6). (D) ChIP-PCR showing the enrichment of H3K27ac at the promoter region of LINC00599 (n=6). Data were expressed as the mean ± SEM. *\*, P<0.05; ***, P<0.001.* SE: super enhancer; E: typical enhancer; Nor: normoxia; Hyp: hypoxia; NC: negative control; si-ZNF263: ZNF263 siRNA.

